# Herpes simplex virus pUL56 abolishes neuronal activity by removing voltage-gated ion channels from the plasma membrane

**DOI:** 10.64898/2026.04.10.717620

**Authors:** Daniel A. Nash, P. Robin Antrobus, Alex S. Nicholson, John Suberu, Martin Potts, Marta A. Almeida, Valeria Lulla, Colin M. Crump, Anton J. Enright, Michael P. Weekes, Janet E. Deane, Stephen C. Graham

## Abstract

Herpes simplex virus 1 (HSV-1) infections of the central nervous system cause encephalitis and are associated with increased risk of neurodegeneration, yet the molecular consequences of lytic infection in human neurones remain incompletely defined. We map the transcriptomic, proteomic and surface-proteome changes induced by HSV-1 across the lytic infection cycle in human iPSC-derived cortical glutamatergic neurones. HSV-1 drives extensive plasma-membrane remodelling, including the removal of voltage-gated sodium, potassium and calcium channels, resulting in a profound loss of synchronous calcium signalling. We identify the viral ubiquitin-ligase adaptor pUL56 as the principal effector of this process: pUL56-dependent depletion of ion channels abolishes coordinated calcium signalling, whereas deletion or mutation of its E3-ligase binding motifs preserves synchrony. Furthermore, expression of pUL56 alone is sufficient to abolish neuronal electrical activity. These findings establish pUL56 as a potent viral suppressor of neuronal excitability and provide potential mechanistic links between HSV-1 infection and neurodegenerative pathology.

## Introduction

Approximately 65% of people under the age of 50 are infected with herpes simplex virus (HSV)-1 (1). HSV-1 infections generally start in peripheral tissues, particularly fibroblasts or keratinocytes of the orofacial or genital mucosa. The virus replicates in these peripheral tissues before infecting sensory or sympathetic neurones, enabling spread to the trigeminal or superior cervical ganglia where they establish a lifelong latent infection (2). The common clinical manifestations of HSV-1 reactivation are cold sores or genital blisters. However, during primary infection or reactivation of latent virus from the trigeminal ganglia, HSV-1 can spread to the central nervous system (CNS) to cause herpes simplex encephalitis (HSE), a necrotizing haemorrhagic inflammatory encephalitis that manifests in the temporal lobes and adjacent limbic tissue (3, 4). HSV-1 infection is the most frequently diagnosed cause of sporadic encephalitis (5), affecting approximately one in 250,000 individuals per year (6, 7). While prompt treatment with the antiviral drug acyclovir reduces mortality rates from 70% to under 20% (6, 8), approximately 50% of HSE survivors experience significant neurological sequelae (9–11). Furthermore, HSV-1 infection is linked to increased risk of Alzheimer’s disease (AD) and related dementias (12–15), and infection studies using human induced pluripotent stem cell (iPSC)-derived 3D models recapitulate multiple AD-like pathologies including amyloid fibrils, neuroinflammation and diminished neural network functionality (16).

Much is known about the molecular consequences of lytic HSV-1 infection in peripheral cells (17–19), and there is substantial literature describing infection plus mechanisms of latency in sensory neurones (20, 21). However, much less is known about the molecular consequences of lytic HSV-1 infection in cells of the human CNS. Animal models enable the study of HSV-1 infection pathology (22), and organoid models represent a promising system for analysing transcriptional changes across different human cell types (23). However, these models are not currently amenable to high resolution proteomic analysis, for example of subcellular protein organisation. Recent advances in iPSC technology allow rapid differentiation of specific neuronal subtypes, including sensory or cortical neurones following expression of NGN3 or NGN2, respectively (24–26). These iPSC systems complement more complex models by allowing highly efficient infection in an isogenic background (27–29), and are scalable to allow unbiased screening approaches like whole-cell and subcellular proteomics (30, 31) and forward genetic screening (24).

Quantitative Temporal Viromics (QTV) is a powerful technique for identifying how viruses remodel the environment of infected cells over the course of an acute infection (32). Coupled with transcriptomic analysis it allows robust identification of cellular proteins that are specifically degraded by virus-encoded proteins in the absence of mRNA degradation, highlighting viral countermeasures against host cell innate or intrinsic immunity (33). QTV analysis of human keratinocytes infected with HSV-1 identified that viral protein pUL56 is an adaptor for cellular ubiquitin E3 ligases and, combined with plasma membrane (PM) proteomics, identified that pUL56 stimulates degradation of the cellular trafficking factor GOPC, leading to the downregulation of the pattern recognition receptor TLR2 from the surface of infected cells (17).

Recently, we established protocols for highly efficient synchronous infection of human iPSC-derived cortical glutamatergic neurones (i3Neurones), a relevant neuronal subtype for studying encephalitis (27). Here we perform QTV on HSV-1 infected i3Neurones. We show that voltage-gated ion channels are lost from the surface of infected neurones, abolishing their electrical activity, and that this remodelling of the plasma membrane proteome is driven by the HSV-1 ubiquitin E3 ligase adaptor pUL56.

## Results

### QTV analysis of HSV-1 infected i3Neurones

The replication kinetics of HSV-1 strain KOS in differentiated i3Neurones was monitored to identify suitable sampling times post-infection for QTV analysis (Fig. 1A). HSV-1 replication was slower than in peripheral cells (17), with progeny virus production commencing by 9 hours post-infection (hpi) and peak titres obtained at 30 hpi, indicating completion of the replication cycle. QTV was thus performed by synchronously infecting i3Neurones (MOI 5) and collecting lysates at 3, 6, 12, 18, and 30 hpi (Fig. 1B). Mock-infected samples were collected at 12 hpi, with additional mock-infected samples collected at 3 and 30 hpi for transcriptomic analyses. Protein and RNA extracts were prepared for proteomic and transcriptomic analysis, with changes to protein abundance quantified using quantitative MS3 mass spectrometry of 18-plex TMT-labelled samples and using Oxford Nanopore sequencing of cDNA to monitor changes in transcript abundance. Parallel infections confirmed near-100% infection efficiency (Fig. S1), with both proteomic (Fig. S2A) and transcriptomic data (Fig. S3) clustering by infection status and time post-infection rather than by biological replicates. The final analysis quantitated 6135 proteins, of which 61 were viral, and 12,214 transcripts, including 72 viral genes, providing a comprehensive view of proteome and transcriptome changes during HSV-1 infection. 1446 of the proteins identified by mass spectrometry were not identified in the previous proteomic study of HSV-1 infected keratinocytes (17) and many are likely to be neurone-specific. Both datasets can be interrogated using an interactive “plotter” worksheet in Dataset S1.

**Figure 1.**
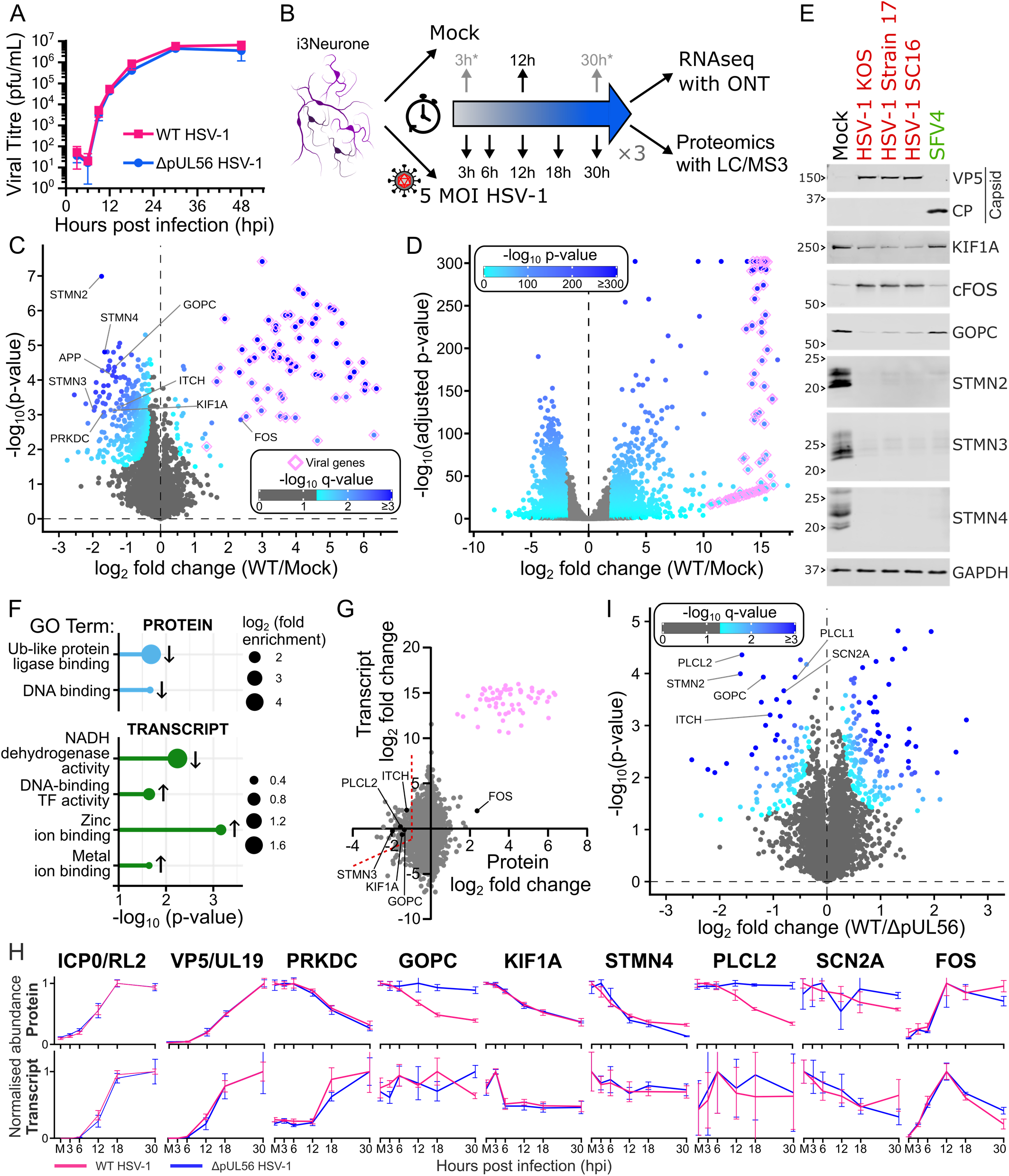
Quantitative Temporal Viromics (QTV) of HSV-1 infection in cortical glutamatergic neurones. (*A*) Single-step (MOI 5) replication curve of wildtype (WT) and ΔpUL56 HSV-1 in day 14 i3Neurones. Mean ± SD for three independent experiments is shown. (*B*) Schematic of the QTV workflow using day 14 i3Neurones. Mock samples were harvested at 12 hpi for all analyses, and additionally at 3 and 30 hpi for transcriptomics (RNAseq) using Oxford Nanopore Technology (ONT). (*C*) Changes in i3Neurone protein abundance at 30 hpi with WT HSV-1. Horizontal axis shows average fold change and vertical axis shows significance (two-sided t-test) for three independent experiments. Proteins are coloured by FDR-corrected significance and viral proteins are outlined in pink. (*D*) Changes in i3Neurone transcript abundance at 30 hpi with WT HSV-1. Horizontal axis shows average fold change and vertical axis shows FDR-adjusted significance for three independent experiments. Significantly altered genes (log_2_ fold change ≥ 2 and p ≤ 0.05) are blue and viral transcripts are outlined in pink. (*E*) Immunoblot analysis of day 14 i3Neurones at 16 hpi (MOI 10) with HSV-1 strains KOS, S17 and SC16, or Semliki Forest virus (SFV)4. Changes in KIF1A, cFOS and GOPC are HSV-1 specific, whereas abundance of stathmins (STMN2–4) is altered in all infections. Capsid proteins VP5 (HSV) and CP (SFV4) confirm successful infection and GAPDH is a loading control. (*F*) Gene Ontology (GO) analysis of proteins (top, blue) and transcripts (bottom, green) with significantly changed abundance at 30 hpi in HSV-versus mock-infected i3Neurones. Significant GO molecular function terms are shown as bubbles sized by magnitude of enrichment, with significance (FDR-adjusted p-value) plotted on the horizontal axis and arrows indicating whether the term was enriched amongst the up- or down-regulated gene products. (*G*) Changes in protein (horizontal) versus transcript (vertical) abundance in mock-versus HSV-infected i3Neurones at 30 hpi. Cellular and viral gene products are grey and pink, respectively, with selected cellular gene products highlighted (black). Potential targets of HSV-1 directed proteasomal degradation (larger fold change in protein vs transcript abundance) are to the left of the dotted red line. (H) Normalised abundance traces of selected proteins (top) and transcripts (bottom) across the time course of infection with WT (pink) or ΔpUL56 (blue) HSV-1. Mock-infected samples (M) are shown as 0 hpi. (*I*) Cellular protein abundance changes in day 14 i3Neurones infected (MOI 5) with WT versus ΔpUL56 i3Neurones. Data are plotted as in (*C*), with horizontal axis showing average mock-corrected fold change for three independent experiments.

As anticipated, the most pronounced changes in proteome composition were observed at 30 hpi (Fig. S4A). HSV-1 infection resulted in an overall decrease in cellular protein abundance (observable as a leftward skew in Fig. 1C, Dataset S2) and a significant decrease in the abundance of known HSV-1 targets like PRKDC and GOPC (Fig. 1C), consistent with previous proteomic analysis of keratinocytes (17). The stathmin family proteins STMN2 and STMN4 that are expressed predominantly in the brain (34, 35) were also highly downregulated. In total, 411 proteins were significantly downregulated by HSV-1 at 30 hpi, with 112 being less than half as abundant as in mock infection, and 86 proteins were significantly upregulated, of which 61 were viral. FOS was the most upregulated cellular protein, likely due to mRNA stabilisation following HSV-1 infection (36, 37). Transcriptomic changes were also most pronounced at 30 hpi (Fig. S4B), with 1715 transcripts significantly downregulated and 1322 (including 72 viral transcripts) significantly upregulated (Fig. 1D, Dataset S3). To validate the proteomics and probe whether abundance changes represent a specific response to HSV-1 infection, immunoblots were performed on lysates prepared from i3Neurones infected for 24 h at MOI 10 with HSV-1 strains KOS, S17 or SC16, or with the neurotropic RNA virus Semliki Forest virus 4 (SFV4). KIF1A and GOPC were degraded only upon infection with HSV-1, whereas stathmin family proteins STMN2, STMN3 and STMN4 were all decreased following infection with either HSV-1 or SFV4, suggesting that their degradation represents a general neuronal response to virus infection (Fig. 1E). Gene Ontology (GO) analysis of gene products significantly changed in abundance at 30 hpi (Fig. 1F) revealed no molecular function terms enriched among upregulated proteins, but ‘ubiquitin-like protein ligase binding’ was enriched among downregulated proteins. Among downregulated transcripts, ‘NADH dehydrogenase activity’ was significantly enriched, and three other terms related to DNA-binding or metal ion-binding proteins were enriched among upregulated transcripts. There was no significant enrichment of interferon-stimulated genes amongst the upregulated proteins or transcripts at any timepoint.

Hijacking the host ubiquitin ligase axis is crucial during HSV-1 infection; viral proteins that possess ubiquitin ligase activity, or that act as ligase adaptor proteins, target host proteins for degradation to counteract innate immune responses (17, 38–41). Proteins specifically targeted for ubiquitination and degradation in i3Neurones could thus represent novel neuronal antiviral restriction factors that HSV-1 can overcome. To identify proteins that are actively degraded, the log_2_ fold change in protein abundance between WT and mock infection was compared with the change in transcript abundance (Fig. 1G, Fig. S5). Proteins such as GOPC and ITCH, which are targeted for ubiquitin-mediated proteasomal degradation by HSV-1 pUL56, were degraded despite their transcript levels remaining steady or increasing. Given the known role of pUL56 target protein GOPC in regulating trafficking of AMPA receptors (42), the contribution of pUL56 to HSV-1 infection of i3Neurones was investigated. The replication kinetics of HSV-1 lacking pUL56 (ΔpUL56) in i3Neurones was indistinguishable from the WT virus (Fig. 1A), as previously observed for keratinocytes (17, 43). QTV of ΔpUL56 HSV-1 infection was thus performed using the same timepoints as for the WT virus. Again, replicates clustered by time post-infection rather than by biological replicates (Figs S2B, S3). Furthermore, the clustering analysis suggests that the transcriptomes of i3Neurones infected with WT and ΔpUL56 HSV-1 are highly similar (Fig. S3), and at 30 hpi there were no robust differences in how either virus alters the transcriptome relative to mock infection (Fig. S6). In contrast, there were significant changes in how WT and ΔpUL56 HSV-1 alter the proteome of infected i3Neurones (Fig. 1H). For example, the abundance of membrane-associated proteins PLCL1, PLCL2 and SCN2A that modulate neuronal signalling (44– 46) were reduced at 30 hpi with WT HSV-1, but significantly rescued following infection with HSV-1 lacking pUL56.

### pUL56 targets voltage-gated ion channels for removal from the plasma membrane

Given the importance of the cell surface in mediating communication between neurones, proteomic analysis was performed to identify whether pUL56 affects the surface abundance of SCN2A or other PM proteins. i3Neurones were synchronously infected (MOI 5) with WT or ΔpUL56 HSV-1, or mock infected, and at 18 hpi cells were surface labelled with biotin before the biotinylated proteins were enriched via streptavidin capture and analysed by mass spectrometry (Fig. 2A). Filtering for proteins with GO cellular compartment annotations “plasma membrane”, “cell surface”, or “extracellular region” resulted in 1451 quantified host proteins, with datasets clustering by infection status rather than biological replicates (Fig. S7). Infection with WT HSV-1 dramatically altered the PM proteome of i3Neurones, with 138 and 270 cell surface proteins significantly up- or down-regulated, respectively (Fig. 2B, Fig. S8, Dataset S4). Strikingly, 28 proteins were less than a quarter as abundant at the surface of infected vs mock-infected neurones. These included voltage-gated sodium, potassium and calcium channel proteins, as well as proteins involved in cell adhesion and receptor transduction, plus the known target of HSV-1 surface downregulation nectin-1 (47). The voltage-gated sodium channel component SCN2A, a target of pUL56-dependent degradation in the whole-cell proteomics, was one of the most highly downregulated surface proteins, suggesting that pUL56 mediates its plasma membrane removal and subsequent degradation. Comparison between WT and ΔpUL56 HSV-1 infections confirmed that pUL56 mediates the downregulation of SCN2A and 13 other voltage-gated ion channel proteins, plus the glutamate-gated ion channel GRIA2 (Fig. 2C). 31 proteins were more than 2-fold upregulated at the plasma membrane of WT HSV-1 versus mock-infected cells, including 7 where the abundance declines back to mock levels in neurones infected with ΔpUL56 HSV-1. Of these seven, two belong to the ANKRD13 family of proteins that mediate the endocytic removal of ubiquitinated plasma membrane proteins from the cell surface (48). Examples of proteins downregulated from the cell surface during infection with both WT and ΔpUL56 HSV-1 (nectin-1), that are downregulated by WT HSV-1 but rescued in ΔpUL56 infection (ion channel proteins), or that are upregulated only during ΔpUL56 HSV-1 infection (ANKRD13 family proteins) are shown in Fig. 2D, and the full PM proteomics dataset can be interrogated using an interactive “plotter” worksheet in Dataset S1. GO analysis of the proteins significantly upregulated in either the WT-versus-mock or ΔpUL56-versus-WT HSV-1 infections did not reveal any significantly enriched molecular function terms. However, amongst significantly downregulated proteins many terms were enriched (Fig. 2E, F), including multiple terms related to voltage-gated ion channel activity. Previous studies have reported downregulation of specific ion channels and/or disrupted electrical activity following HSV-1 infection of sensory or cortical neurones (49–57). However, the full extent of ion channel modulation and the underlying mechanism was unclear. The significant changes in PM ion channel abundance between WT and ΔpUL56 HSV-1 infected neurones strongly suggests that pUL56 alters ion flux to disrupt neuronal electrical activity.

**Figure 2.**
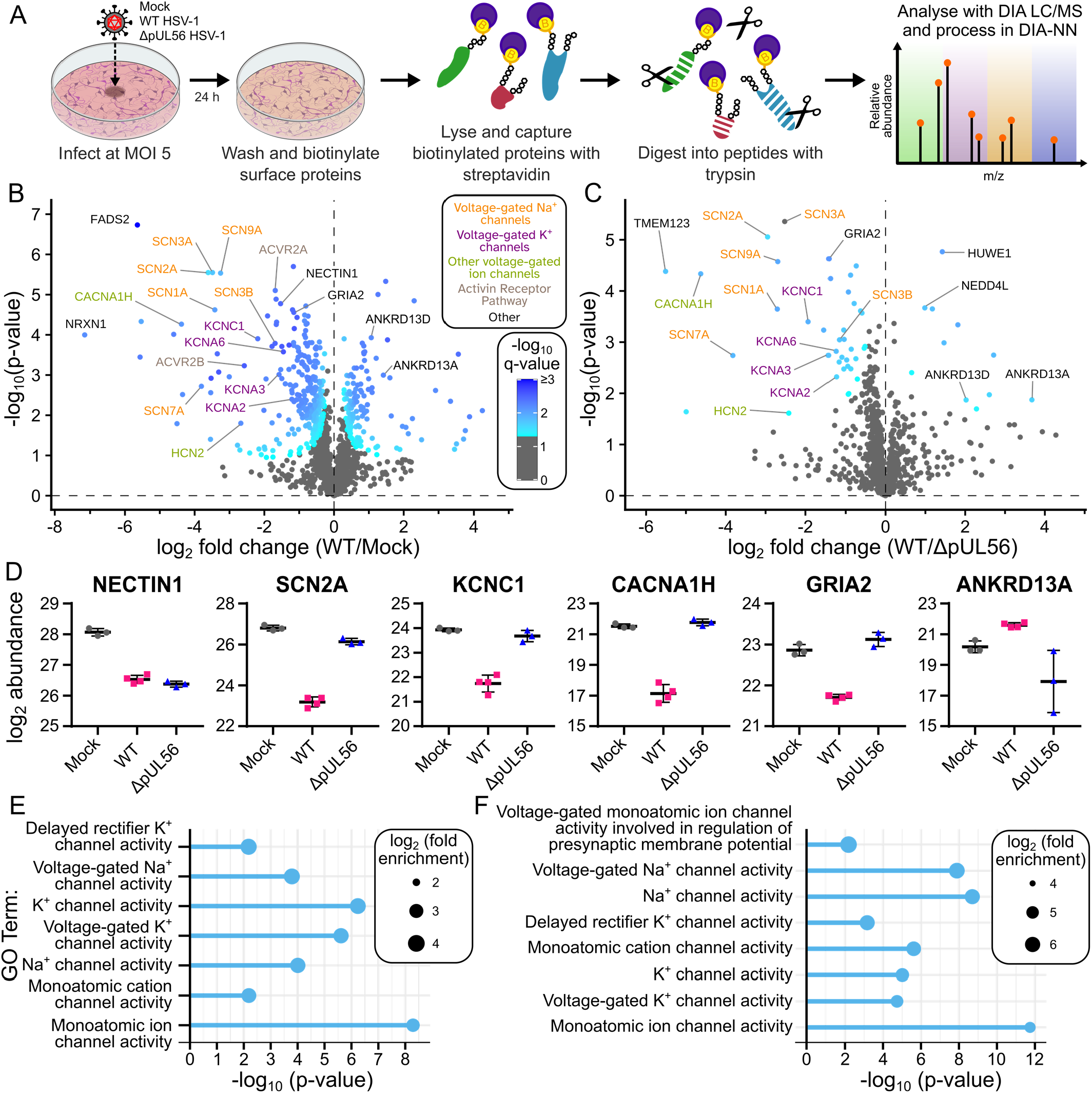
pUL56 reduces the plasma membrane abundance of voltage-gated ion channels. (*A*) Schematic of the plasma membrane proteomics (PMP) workflow using day 14 i3Neurones. (*B* and *C*) Change in PM protein abundance at 24 hpi, showing (*B*) WT versus mock-infection, or (*C*) WT versus ΔpUL56 HSV-1 infection. Horizontal axis shows average fold change and vertical axis shows significance (two-sided t-test) for three independent experiments. Proteins are coloured by FDR-corrected significance and selected proteins with significantly altered abundance are labelled. (*D* and *E*) Enrichment of GO molecular function terms amongst proteins significantly downregulated from the PM in (D) WT versus mock-infection, or (*C*) WT versus ΔpUL56 HSV-1 infection. Bubble sizes represent magnitude of enrichment and significance (FDR-adjusted p-value) is plotted on the horizontal axis. (*F*) PM abundance of HSV-1 entry receptor NECTIN1, voltage-gated Na^+^ (SCN2A), K^+^ (KCNC1) and Ca^2+^ (CACNA1H) ion channels, glutamate receptor subunit 2 (GRIA2), and ubiquitin reader protein ANKRD13A in mock-infected versus WT or ΔpUL56 HSV-1 infected i3Neurones. Individual data points and mean abundance ± SD are shown.

### pUL56 silences calcium signalling during infection via an interaction with E3 ubiquitin ligases

Like many neurotropic viruses, HSV-1 infection alters neuronal electrophysiology (49, 50, 58–63) but the mechanism by which it does so was unknown. To investigate the consequence of pUL56 altering the abundance of voltage-gated ion channels at the neuronal surface, changes to calcium signalling during infection were analysed as a proxy for neuronal electrical activity (64). i3Neurones are electrically mature after approximately 30 days (30), so i3Neurones that had been transduced with a fluorescent calcium biosensor were synchronously infected (MOI 5) on day 42 with WT or ΔpUL56 HSV-1, or mock infected. To investigate whether binding of pUL56 to NEDD4-family E3 ligases is required for any changes in electrical activity, neurones were also infected with pUL56 AAXA123 HSV-1 where all three PPXY motifs had been mutated to reduce E3 ligase binding affinity (17). Preliminary experiments showed that repeated calcium-flux kinetic measurements require *de novo* protein expression to replace biosensor molecules that bleach during the fluorescent imaging, and that by 24 hpi with HSV-1 this fluorescence recovery is inhibited, presumably by host shutoff. Therefore, calcium imaging was conducted at 12, 24, and 48 hpi on separate wells of i3Neurones (Fig. 3A). At 50 hpi the wells measured at 12 hpi were treated for 1 h with 1 μM tetrodotoxin (TTX) to eliminate electrical activity via inhibiting sodium channels before being re-imaged. Immunostaining of the wells measured at 24 hpi confirmed near-100% infection (Fig. S9). Calcium ion flux remains unaffected by HSV-1 infection at 12 hpi (Fig. 3B and C, Movie S1). However, by 24 hpi synchronous firing is entirely abolished in cells infected with WT HSV-1, while it persists at levels comparable to mock-infected samples in neurones infected with either ΔpUL56 or pUL56 AAXA123 HSV-1 (Fig. 3B, Movie S2). The number of active neurones declines in all infections, but WT HSV-1 causes a substantially greater reduction at 24 hpi and near-complete cessation of neuronal activity by 48 hpi (Fig. 3C, Movie S3). In contrast, approximately two-thirds of the cells infected with ΔpUL56 or pUL56 AAXA123 HSV-1 remain active at 48 hpi. Representative traces of calcium biosensor intensity at 48 hpi illustrate how mock-infected neurones and those infected with ΔpUL56 or pUL56 AAXA123 HSV-1 exhibit synchronous intracellular calcium bursts, albeit with a reduced peak amplitude in the infected neurones, whereas neurones infected with WT HSV-1 have no synchronous activity, akin to treatment of neurones with TTX (Fig. 3D). pUL56 is thus required for the dramatic inhibition of neuronal calcium signalling in infected neurones, and this inhibition requires the PPXY motifs of pUL56 that mediate association with ubiquitin E3 ligases.

**Figure 3.**
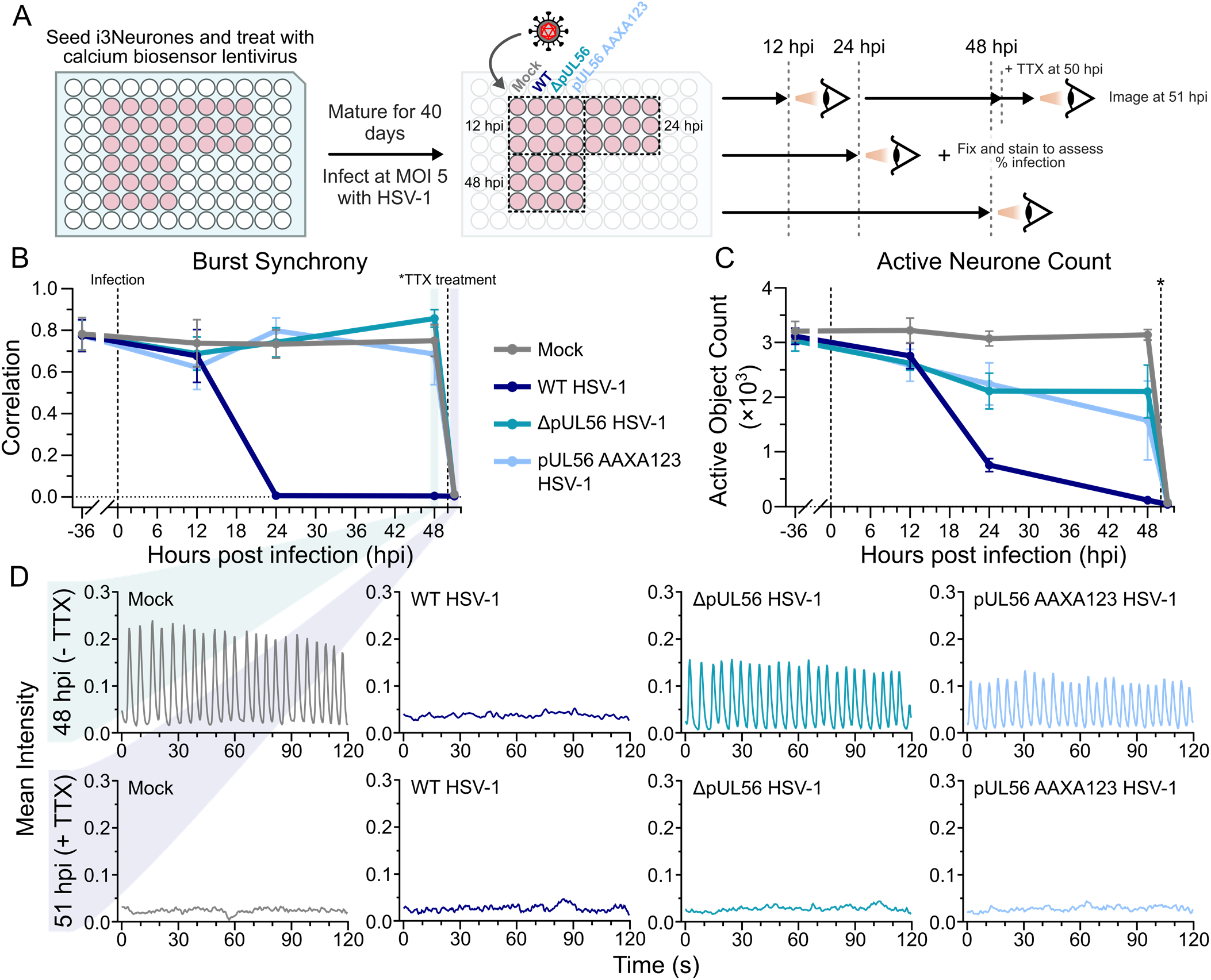
HSV-1 infection disrupts calcium signalling in i3Neurones. (*A*) Schematic diagram of the experimental design. Calcium biosensor transduced day 40 i3Neurones were infected with WT, ΔpUL56 or pUL56 AAXA123 HSV-1 (MOI 5), or mock-infected, and imaged at 12, 24 or 48 hpi. Cells imaged at 12 hpi were treated with 1 μM tetrodotoxin (TTX) at 50 hpi and imaged 1 h later. (*B*) WT HSV-1 infection causes a loss of synchronous calcium bursts by 24 hpi, equivalent to TTX treatment, whereas burst synchrony is not reduced in ΔpUL56 and pUL56 AAXA123 infected cells up to 48 hpi. Mean ± SD of three independent experiments performed in technical triplicate are shown. (*C*) The number of actively signalling neurones steadily decreases following ΔpUL56 and pUL56 AAXA123 HSV-1 infection, whereas it drops dramatically at 24 hpi with WT HSV-1. (*D*) Representative plots of mean calcium biosensor signal versus time, showing complete loss of calcium signalling at 48 hpi with WT HSV-1, or at 51 hpi following 1 h treatment of i3Neurones with TTX, but not in cells infected with ΔpUL56 or pUL56 AAXA123 HSV-1 at 48 hpi.

### pUL56 alone is sufficient to abolish neuronal electrical activity

Altered neuronal activity during HSV-1 infection could result from direct pUL56-stimulated downregulation of voltage-gated ion channels, from mistrafficking of these channels as a consequence of pUL56-stimulated GOPC degradation, or via pUL56 directing the activity of a different viral effector protein. To differentiate between these, i3Neurone cell lines were generated where either GOPC expression was knocked down or pUL56 alone was expressed. The catalytically dead Cas9 (dCas9) integrated in the i3Neurone iPSCs was exploited for CRISPR interference (CRISPRi)-mediated GOPC knockdown, via transduction of two distinct CRISPRi guides targeting early exons of GOPC to generate two distinct cell lines (GOPC KD1 and KD2). An established i3Neurone cell line containing a scrambled (SCRM) non-targeting guide line was used as a control (30). Stable genomic integration of expression cassettes for pUL56 or pUL56 AAXA123, both with N-terminal HA epitope tags to aid detection, was achieved via co-transfection using a piggyBac transposase (65, 66). Quantitative PCR of i3Neurones at day 42 confirms successful GOPC knockdown (Fig. 4A) and immunoblot analysis confirms both the absence of GOPC and the expression of pUL56 (Fig 4B). As expected, the abundance of GOPC is reduced in i3Neurones expressing HA-pUL56 but not in cells expressing HA-pUL56 AAXA123 (17).

**Figure 4.**
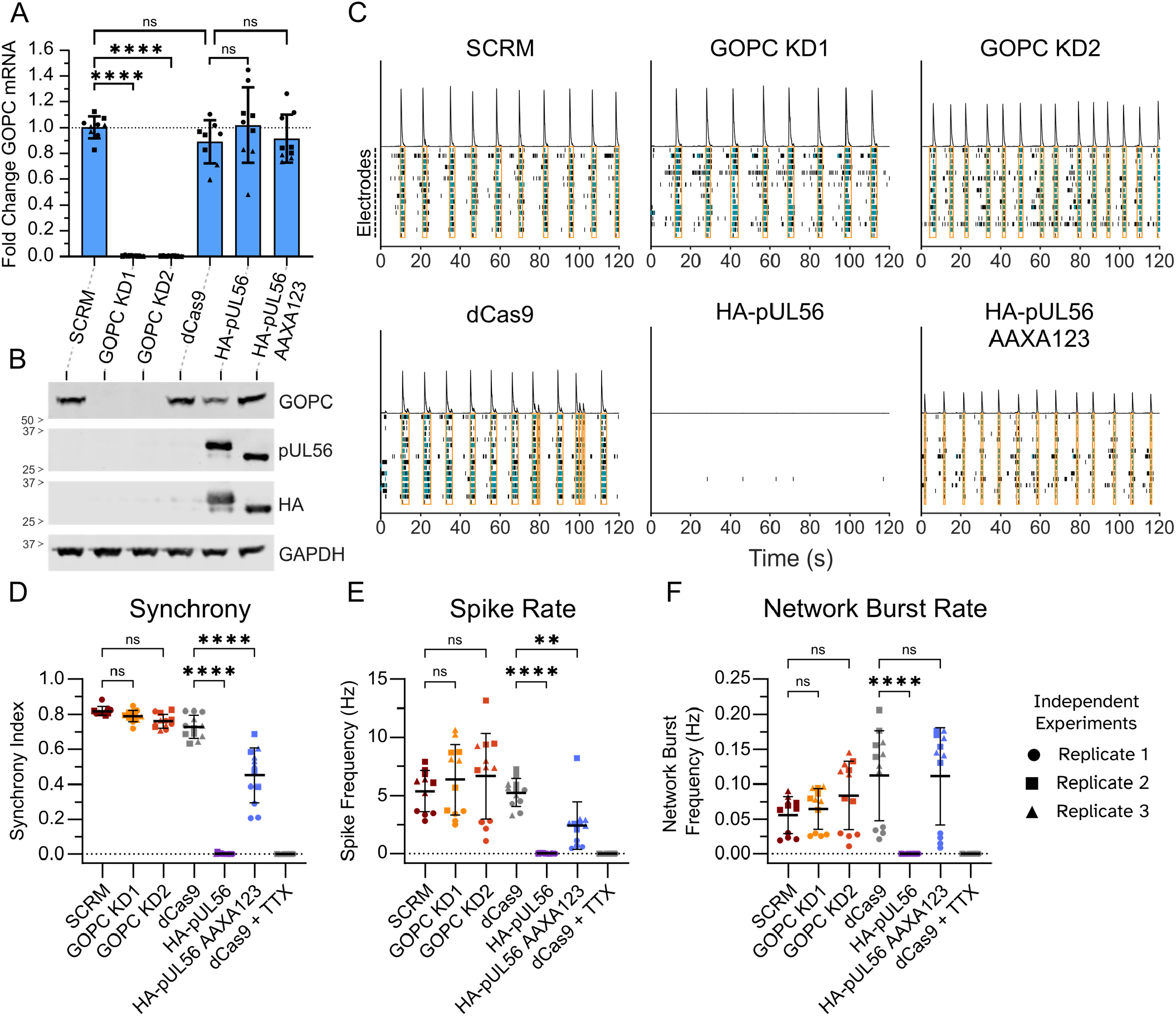
pUL56 halts electrical signalling through a GOPC-independent mechanism. (*A* and *B*) Validation of day 42 i3Neurones expressing catalytically-dead (d)Cas9 that have been stably transduced with guides targeting GOPC (GOPC KD1 and KD2), or a non-targeting control (SCRM), or that stably overexpress HA-tagged pUL56 or pUL56 AAXA123. (*A*) Fold-change in GOPC expression, as assessed by qPCR. Mean ± SD of three independent experiments, each in technical triplicate, are shown as circles, squares and triangles. ns, no significance; ****, p < 0.0001; one-way ANOVA with Sidak’s multiple comparisons test. (*B*) Immunoblot showing pUL56 expression in HA-pUL56 and HA-pUL56 AAXA i3Neurones, and loss of GOPC signal in GOPC KD1 and KD2. Compared to parental cells (dCas9), GOPC abundance is reduced in HA-pUL56 but not HA-pUL56 AAXA i3Neurones. (*C*) Representative multi-electrode array (MEA) raster plots of i3Neurone electrical activity at day 42. For each cell line, frequency histograms of electrical spikes are shown (line trace, *top*), along with individual spikes (black) and bursts (> 5 spikes within 100 ms, cyan) for each electrode (*below*). Network bursts, defined as simultaneous bursts on >35% of electrodes in the well, are boxed in orange. (*D* to *F*) Quantitation of (*D*) spike synchrony, (*E*) spike rate, and (*F*) network burst rate from MEA recordings at day 42. HA-pUL56 expression severely impairs electrical activity, which is partially rescued in the HA-pUL56 AAXA123 line. For the parental (dCas9) line, activity was recorded before and after 1 h incubation with 1 μM TTX. Mean ± SD and individual data points from three biological replicates, each performed in technical quadruplicate, are shown as circles, squares and triangles. ns, no significance; **, p< 0.0001; ****, p < 0.0001; paired measurements one-way ANOVA with Sidak’s multiple comparisons test.

The electrical activity of GOPC knockdown and pUL56-expressing i3Neurones was examined using multi-electrode arrays (MEA). Electrical activity was recorded on day 42 and is shown in Fig. 4C as raster plots for each cell line, illustrating electrical activity as spikes (black line), bursts (green, >5 spikes measured on an electrode within 100 ms), and network bursts (orange box, bursts detected on >35% of electrodes), plus a spike frequency histogram.

Electrically healthy i3Neurones evolve regular, high-frequency spike trains that produce numerous bursts and network bursts (Fig. 4C, SCRM and dCas9). Neither of the GOPC knockdown cell lines had reduced electrical activity compared to the SCRM control, whereas the expression of HA-pUL56 almost completely abolished spike activity. The HA-pUL56 AAXA123 cell line had an intermediate phenotype, with electrical activity decreased but still evident. The firing synchrony (Fig. 4D), spike rate (Fig. 4E), and network burst rate (Fig. 4F) of each cell line were quantified, alongside recordings of the dCas9 neurones following treatment with TTX. There was an absence of synchronous firing in i3Neurones expressing HA-pUL56, similar to cells treated with TTX, whereas synchrony was partially restored in cells expressing HA-pUL56 AAXA123. The spike rate was similarly affected, with no activity observed in HA-pUL56 i3Neurones but some rescue of activity in HA-pUL56 AAXA123 i3Neurones. The network burst frequency of HA-pUL56 AAXA123 i3Neurones was similar to the parental (dCas9) cells, as opposed to HA-pUL56 i3Neurones where no network bursts were observed. Collectively, these data demonstrate that pUL56 expression is sufficient to abolish electrical activity in neurones, and the partial rescue observed in the HA-pUL56 AAXA123 expressing cells confirms that neuronal silencing depends on the ability of pUL56 to efficiently bind ubiquitin E3 ligases. Relative to SCRM i3Neurones, neither GOPC KD1 nor KD2 significantly altered neuronal electrical activity, implying that GOPC does not contribute to the removal of voltage-gated ion channels from the plasma membrane.

## Discussion

This study identifies pUL56 as a major effector of plasma membrane remodelling during HSV-1 infection of human cortical neurones. We demonstrate that pUL56 mediates the removal of voltage-gated ion channels from the neuronal plasma membrane, thereby suppressing electrical signalling during infection. Infection-induced inhibition of calcium ion flux is reversed when the PPXY E3 ligase binding motifs of pUL56 are mutated and, importantly, pUL56 does not require any other viral proteins to abolish neuronal electrical function. Our findings therefore establish that pUL56 is both necessary and sufficient to silence the electrical activity of infected neurones.

While HSV-1 induced substantial proteomic and transcriptomic changes in infected neurones, it did not elicit an interferon-based antiviral response as there was no coordinated upregulation of antiviral genes. This is consistent with our previous observation that HSV-1 infection of i3Neurones is not self-limiting (27). However, there was a rapid decrease in the abundance of several proteins implicated in microtubule assembly and trafficking, including multiple stathmin family members and the kinesin-3 family protein KIF1A (35, 67–69). These effects are not caused by cell death at 30 hpi, as i3Neurones persist for over one week following infection with HSV-1 (27) and cells infected with ΔpUL56 or pUL56 AAXA123 HSV-1 retain synchronous calcium ion flux for at least 48 hpi (Fig. 3). Our results align with similar experiments performed in cerebral organoids, where STMN2 is also heavily downregulated at both the protein and transcription level following HSV-1 infection (23), but we show that degradation of the stathmin-family proteins also occurs in response to SFV4 infection (Fig. 1E). These changes could thus represent a general but concerted antiviral or injury response. Inhibiting axonal transport would directly impact virus particle movement and it would potentiate programmed axonal degradation (70), both acting to impair virus spread. Low STMN2 abundance in motor neurones caused by loss of TDP-43 nuclear localisation is a hallmark of amyotrophic lateral sclerosis (ALS), and the loss or depletion of STMN2 is sufficient to drive ALS-like pathology in mice (71–73). A recent preprint confirms that HSV-1 seropositivity is significantly associated with increased ALS risk (74). While this study implicates the viral E3 ligase ICP0 in driving TDP-43 mislocalisation and aggregation, STMN2 abundance was not investigated. Whether STMN2 degradation is a consequence of TDP-43 mislocalisation or represents an independent driver of ALS in the context of HSV-1 infection remains to be determined.

The observed downregulation of KIF1A was specific to HSV-1 infection. While KIF1A has previously been shown to bind HSV-2 pUL56 (75), the degradation we observe is not induced by HSV-1 pUL56 since KIF1A is lost at an equivalent rate in WT or ΔpUL56 infected neurones (Fig. 1H). KIF1A is a subunit of the kinesin-3 motor protein that mediates anterograde transport of cargo in neurones (69), including nascent particles of HSV-1 (76) and the related alphaherpesvirus pseudorabies virus (77). A direct interaction of KIF1A with pseudorabies virus pUS9 stimulates both virion axonal transport and KIF1A degradation (77), with this degradation suggested to occur at synaptic termini after cargo delivery (78, 79). pUS9 is not expressed in HSV-1 strain KOS used in our study (80), but pUS9 is not absolutely required for HSV-1 axonal transport (81). The HSV-1 specific KIF1A degradation we observe is thus likely to be a result of degradation following synaptic delivery of nascent virions and other cargo, plus lowered KIF1A mRNA abundance following virus infection (Fig. 1H).

For many genes the transcript and protein abundance changes observed during infection differ (Fig. 1G), highlighting extensive post-transcriptional changes during HSV-1 infection. Of the 28 proteins with significantly altered abundance identified in a previous proteomic study of HSV-1 infection using mixed mouse primary cortical cells (82), the only common change we observed was significant downregulation of the Kelch-like ubiquitin E3 ligase KLHL9. Ubiquitin-ligase binding functions are enriched amongst proteins significantly downregulated at 30 hpi (Fig. 1F), consistent with previous reports of HSV-1 hijacking host ubiquitin pathways to eliminate antiviral factors and reshape the infected cell environment (17). pUL56 has previously been identified as an adaptor for the ubiquitin-proteasome axis, binding to E3 ubiquitin ligases to target proteins such as GOPC for degradation (17). The loss of ion channels from the plasma membrane is rescued in ΔpUL56 infection and calcium ion flux is restored either when pUL56 is absent during infection or when its PPXY motifs are mutated, highlighting the requirement for an interaction between pUL56 and NEDD4-family E3 ligases to mediate this process. Furthermore, the loss of synchrony mimics TTX treatment, strongly indicating that suppression of neuronal activity arises from the loss of cell surface voltage-gated ion channel activity.

We demonstrate that pUL56 expression alone is sufficient to abolish neuronal electrical activity. MEA recordings show that pUL56, in the absence of any other viral gene, eliminates spikes, bursts, and network-level synchrony, while mutation of the pUL56 PPXY motifs produces a partial rescue. In contrast, GOPC knockdown has no impact on electrical activity. This is similar to the observation in HEK293T cells that pUL56 mediates degradation of volume-regulated anion channels in an E3-ligase dependent but GOPC-independent manner (83). In i3Neurones, we propose that pUL56 colocalises with voltage-gated ion channels to cause their ubiquitination and subsequent removal from the plasma membrane. The plasma membrane localisation of most of the targeted ion channels is controlled by ubiquitination by NEDD4-family E3 ligases (84–89), the same family that pUL56 binds (17, 90–92). Furthermore, the increased surface abundance of E3 ligases NEDD4L and HUWE1 during WT infection suggests recruitment by pUL56. Interestingly, two members of the ANKRD13 family of ubiquitin-reader proteins are also upregulated at the plasma membrane during WT infection, consistent with their recruitment either by pUL56 directly or by targets of pUL56-directed ubiquitination. These proteins form complexes that specifically bind to lysine-63-linked polyubiquitin, a form preferred by NEDD4 ligases (93, 94), on surface proteins, causing their endocytic internalisation and lysosomal degradation (48). This evidence supports a model in which pUL56 recruits E3 ligases to the plasma membrane to ubiquitinate voltage-gated ion channels, likely via K63 polyubiquitination, and then recruits ANKRD13 proteins to mediate their removal from the neuronal surface.

The vast majority of adults worldwide are infected with HSV-1 (1). Although it remains unclear whether the virus can establish latent infections in human cortical neurones, there is extensive evidence of HSV-1 DNA in the brain of individuals that have not experienced clinical HSE (95, 96). These observations are consistent with sporadic sub-clinical infection of the CNS that arise following reactivation of latent virus in the trigeminal ganglia and are either restricted by intrinsic immune factors (97) or controlled via innate immune sensing (98, 99). Our data suggest that during (subclinical) lytic infection the affected neurones will be electrically silenced within 24 h. Although it remains unclear whether disruption of neuronal firing is advantageous for virus replication and spread, disruption of electrical activity is clearly detrimental for brain function and represents a major contributing factor to many neurological and degenerative disorders, including AD (100–103). Indeed, a recent study demonstrated that subclinical neonatal infection with HSV-1 leads to persistent CNS infection and poor cognitive performance of adult mice (104). Epidemiological studies have linked HSV-1 seropositivity to a higher risk of dementia (105–107) and suggested that HSV antiviral treatment may reduce dementia risk (108), although not all studies confirm this association (109). Treatment with the acyclovir prodrug valcyclovir improved memory and cognitive function in HSV seropositive schizophrenia patients (110), but a recent randomised controlled trial demonstrated no effect of valcyclovir on cognitive worsening of HSV seropositive patients with early symptomatic AD (111). While neuronal silencing by pUL56 during subclinical CNS infections provides a compelling molecular mechanism that could link HSV-1 infection with cognitive impairment, the extent to which HSV-1 infection (and thus pUL56 expression) predisposes individuals to AD or alters its preclinical progression remains to be confirmed.

Our study defines pUL56 as a potent viral suppressor of neuronal excitability, acting through ubiquitin-dependent removal of voltage-gated ion channels from the neuronal surface. The discovery that a single viral protein can shut down neuronal networks provides new insight into how neurotropic viruses manipulate the nervous system. There is increasing interest in the use of non-replicative (nr)HSV-1 as a neurotropic gene delivery vector (112). While modern nrHSV-1 vectors show very low levels of viral gene expression (113), there have been reports of virion-delivered proteins altering neuronal activity (114). Given that pUL56 is packaged into HSV-1 virions (115), the safety of nrHSV-1 vectors might be further enhanced by removal of the UL56 gene.

## Materials and Methods

### Cell culture

All cells were incubated in a humidified 5% CO_2_ atmosphere at 37°C and regularly tested to confirm the absence of mycoplasma. iPSCs and i3Neurones were maintained and differentiated as detailed in (27). Briefly, iPSCs containing a doxycycline-inducible Ngn2 transcription factor (25) were maintained in Essential 8 medium (Gibco) on Matrigel (Corning) coated plates. Differentiation was initiated by addition of 2 μg/mL doxycycline in Dulbecco’s modified Eagle Medium (DMEM)/F-12 (Gibco) supplemented with 1× N2 (Gibco), 1× non-essential amino acids (NEAA, Gibco) and 2 mM L-glutamine (Gibco). For the first day of differentiation, the medium was supplemented with 50 nM Chroman 1 (Bio-Techne) and the medium was changed daily. On day 3, cells were dissociated with StemPro Accutase (Gibco) and frozen at -70°C in 90% v/v KnockOut Serum Replacement (Gibco) plus 10% v/v DMSO. Thawed day 3 i3Neurone precursors were matured in cortical neurone (CN) medium, comprising Neurobasal Plus Medium (Gibco) supplemented with 1× B27 (Gibco), 10 ng/mL Brain Derived Neurotrophic Factor (BDNF, PeproTech), 10 ng/mL Neurotrophin-3 (NT-3, PeproTech) and 1 μg/mL laminin (Gibco), on surfaces pre-coated with 100 μg/mL poly-L-ornithine (PLO). CN medium was supplemented with 2 μg/mL doxycycline only when seeding the cells. Half media changes were performed twice weekly until day 14, then thrice weekly. Vero and HEK293T cells were grown in DMEM with 2 mM L-glutamine and 10% v/v foetal bovine serum (complete DMEM). BHK21 cells were grown in DMEM with 2 mM L-glutamine, 5% v/v foetal bovine serum, 0.3% tryptose phosphate broth, 100 U/mL penicillin and 100 μg/mL streptomycin.

### GOPC CRISPRi knockdown

CRISPRi knockdown cell lines were generated via lentivirus transduction. Guide RNAs targeting GOPC were designed using the CRISPick server (Broad Institute) to target exon1. Primers encoding the guides (Table S1) with complementary overhangs were cloned into the BpiI restriction site of the plasmid pKLV. Lentiviruses were generated and iPSCs were transduced as described in (30). After one week of puromycin selection, the GOPC knockdown i3Neurones were differentiated as described above.

### pUL56 overexpression

N-terminally HA-tagged pUL56 was amplified by PCR (Table S1) and cloned via NEBuilder HiFi DNA assembly (New England Biolabs) into vector iC10, a kind gift from Matthew Nelson and Michael Ward (National Institutes of Health). HA-tagged pUL56-AAXA (17) was subcloned into this vector following restriction digest using NotI and XhoI sites introduced when cloning HA-pUL56. iPSCs were co-transfected with a 1:3 ratio of the K13 piggyBac transposase plasmid, a kind gift from Matthew Nelson and Michael Ward (National Institutes of Health), and the relevant expression plasmid using Lipofectamine Stem (Invitrogen). Two days post-transfection, 125 μg/mL hygromycin B was added to the E8 medium and selection was maintained for 1 week before surviving cells were differentiated as described above.

### Viruses

Stocks of WT, ΔpUL56 and pUL56 AAXA HSV-1 strain KOS that had been reconstituted from bacterial artificial chromosomes (17) were grown on Vero cells, as were HSV-1 strains 17 and SC16, and all were filter-purified from the supernatant of Vero cells treated with NaCl and dextran sulfate as described in (27). SFV4 was rescued from the plasmid pSP6-SFV4 (116) in BHK21 cells and further amplified using MOI 0.01 infection. Media-derived virus was clarified using 0.4 μm filter and concentrated using Amicon Ultra centrifugal filters with 100 kDa molecular weight cut-off. The resulting virus was titrated on BHK21 by plaque assay (117).

### Single-step growth curves

Day 14 i3Neurones grown in 24-well plates were infected (MOI 5) for 1 h with WT or ΔpUL56 HSV-1, incubated for 1 min with citric acid and washed three times with phosphate buffered saline (PBS) before being overlaid with conditioned medium as described in (27). At 3, 6, 9, 12, 18, 30 and 48 hpi, neurones were harvested by freezing-thawing and virus titres were assessed by plaque assay on Vero cells (27).

### Immunoblotting

Day 14 i3Neurones were infected (MOI 10) for 1 h with HSV-1 strains KOS, S17 and SC16, or SFV4 as described in (27). At 16 hpi, cell lysates were prepared by detaching neuronal cell layers with ice-cold PBS containing 1% v/v protease-inhibitor cocktail, centrifugation to collect cells and lysis in radioimmunoprecipitation assay (RIPA) buffer as described in (27). Following separation by SDS-PAGE, proteins were transferred onto 0.45 μm Protran nitrocellulose membranes (Cytivia) using the Mini-PROTEAN system (Bio-Rad) and blocked in TBS (50 mM Tris pH 7.6, 150 mM NaCl) supplemented with 5% w/v skim milk powder (Marvel). Primary and secondary antibody incubations were performed in TBS supplemented with 0.1% TWEEN-20 and 5% (w/v) skim milk powder using the following antibodies: HSV-1 VP5, (118); SV4 CP, kindly provided by Andres Merits (119); KIF1A, BD Transduction Laboratories 612094; cFOS, Cell Signalling Technologies 2250T; GOPC, Abcam Ab133472; STMN2, Novus Biologicals NBP1-49461; STMN3, ProteinTech 11311-1-AP; STMN4, ProteinTech 12027-1-AP; GAPDH, GeneTex GTX28245; HSV-1 pUL56, (17); HA, Cell Signalling Technologies C29F4; Anti-rabbit 800, LI-COR 926-32213; Anti-rabbit 680, LI-COR 926-68023; Anti-mouse 800, LI-COR 926-32210; Anti-mouse 680, LI-COR 926-68020. Immunoblots were imaged using an Odyssey CLx (LI-COR) and analysed using Image Studio Lite (LI-COR).

### Quantitative PCR

Cells were lysed, RNA extracted using the PureLink RNA Mini extraction kit (Invitrogen), cDNA generated, and qPCR was performed using TaqMan master mix and FAM-labelled assay probes (Bio-Rad) for GAPDH (Hs02786624_g1) and GOPC (Hs01100473_m1) as described in (27). Expression changes were measured using the ΔΔCt method (120).

### Quantitative temporal proteomics

3×10^5^ day 3 i3Neurones were seeded in PLO-treated 24-well plates (2 wells per condition per timepoint per replicate, plus one additional well per infection for microscopy) and matured to day 14 before being synchronously infected (MOI 5) with WT or ΔpUL56 HSV1, or mock-infected. Inoculum was removed after 1 h and cells were washed thrice with PBS before being overlaid with conditioned media. At 3, 6, 12, 18 and 30 hpi, medium was removed and cells were washed once with PBS, which was subsequently aspirated and plates frozen at -70°C. For lysis, plates were thawed and cells scraped into 75 μL of lysis buffer (6 M guanidine HCl, 50 mM HEPES pH 8.5). Lysates from duplicate wells were pooled, snap-frozen in liquid nitrogen and stored at -70°C. At 18 hpi, the additional wells were fixed and stained for ICP4 to assess percentage of infection as described in (27). Lysates were thawed, vortexed extensively, sonicated at maximum power for 5 cycles of 30 s and clarified twice by centrifugation (13,000 × g, 10 min, 4°C). 150 μL of supernatant was reduced (5 mM dithiothreitol, 20 min) then alkylated (14 mM iodoacetamide, 20 min) at room temperature (RT), protected from light, before quenching with excess DTT. Samples were diluted four-fold with 200 mM HEPES pH 8.5 before being supplemented with 3 μL of LysC protease (Promega) for 3 h at RT. The LysC-digested sample was diluted three-fold with 200 mM HEPES pH 8.5 before addition of 1.5 μg of sequencing-grade trypsin (Promega) and incubation overnight at 37°C on a shaking incubator. The reaction was quenched by addition of 5% v/v formic acid and clarified by centrifugation (21,000 × g, 10 min). Digested peptides were desalted (Pierce Peptide Desalting Spin Columns, ThermoFisher), eluted in 50% v/v acetonitrile plus 0.1% trifluoroacetic acid, dried (45°C, 1 Torr) and resuspended in 20 μL of 200 mM HEPES pH 8.5. The concentration of resuspended peptides was analysed using the Pierce Quantitative Colorimetric Peptide Assay (ThermoFisher) as per the manufacturer’s instructions.

30 μg of peptides per sample was diluted to a total volume of 14 μL with 200 mM HEPES pH 8.5, supplemented with 100 µg of TMTpro 18-plex reagents (ThermoFisher) in anhydrous acetonitrile (Acros Organics), vortexed and then incubated at RT for 1 h. The labelling reaction was quenched with 5% v/v hydroxylamine (Sigma Aldrich) for 15 min at RT then formic acid was added to 1% v/v. Sample labelling was as indicated in Table S2. All 18 TMT-labelled samples were combined at a 1:1 ratio, desalted and analysed by single injection on an Orbitrap Fusion Lumos (ThermoFisher) to estimate label contribution. Samples were combined based on observed reporter ion abundances from the single injection to achieve equal label loading, desalted, and fractionated by high-pH reverse-phase chromatography using an Ultimate 3000 ultra-high-performance chromatography (UHPLC) system (Thermo Scientific) equipped with a Kinetex Evo C18 (1.7 μm, 2.1 x 150 mm) column (Phenomenex). The mobile phase used was: HPLC-grade water 3:97 v/v (solvent A); 100% v/v LCMS-grade acetonitrile (solvent B) and 200 mM ammonium formate in HPLC-grade water, pH 10 (solvent C). Eluent C was maintained at 10%, while A and B were altered over the fractionation gradient. The elution gradient was 0–19% B in 10 min; 19–34% B in 14.25 min and 34–50% B in 8.75 min, followed by a 10 min wash of 90% B. Flow was set at 0.20 mL/min for the first 5 min while peptides, which were reconstituted in 40 μL of eluent C, were loaded onto the column. Thereafter, the flow rate was increased to 0.40 mL/min for the rest of the protocol. 168 fractions were collected every 0.25 min between 10 and 52 min of the fractionation run. The fractions were concatenated into 12 pools which were dried in a refrigerated vacuum centrifuge.

Mass spectrometry data were acquired using an Orbitrap Fusion Lumos (Thermo Scientific) equipped with an EASYspray source and coupled to an RSLC3000 nano UPLC (ThermoFisher Scientific). Peptides were fractionated using a 50 cm C18 PepMap EASYspray column maintained at 40°C with a solvent flow rate of 300 nL/min. A gradient was formed using solvent A (0.1% formic acid) and solvent B (80% v/v acetonitrile, 0.1% formic acid) rising from 3 to 7% solvent B by 7 min, from 7 to 37% solvent B by 180 min followed by a 4 min wash at 95% solvent B. MS acquisition used a MultiNotch MS3-based TMT method using the following settings: MS1, 380-1,500Th, 120,000 resolution, 2.0×10^5^ automatic gain control (AGC) target, 50 ms maximum injection time; MS2, quadrupole isolation at an isolation width of m/z 0.7, collision-induced dissociation (CID) fragmentation (normalised collision energy (NCE) 34) with ion-trap scanning in turbo mode with 1.5×10^4^ AGC target; MS3, in synchronous precursor selection mode the top ten MS2 ions were selected for HCD fragmentation (NCE 45) and scanned in the Orbitrap at a resolution of 60,000 with an AGC target of 1.0×10^5^ and a maximum accumulation time of 150 ms. Ions were not accumulated for all parallelizable time. The entire MS/MS/MS cycle had a target time of 3 s. Dynamic exclusion was set to ±10 p.p.m. for 70 s. MS2 fragmentation was triggered on precursors of 5.0×10^3^ counts and above.

Raw mass spectra were processed using MaxQuant version 2.5.2.0 (121, 122) searching against a manually curated HSV-1 strain KOS proteome and the human proteome (UP000005640, accessed on 17.04.2024). Reporter ion intensities were corrected using lot-specific isotopic impurity values (Table S3). Default options within MaxQuant were used with match between runs analysis, and QTV whole cell lysate data (Fractions 1-14) from (17) were included as a separate parameter group to improve search power. Data filtering and label normalisation were performed in R (Software S1). All potential false positives (contaminants, reverse hits, proteins identified only by site) were removed. Scale factors for each label were calculated by comparing the column sum of all proteins within a label to the median column sums of all labels. Each reporter ion intensity was scaled for normalisation and a log_2_ transformation was applied for downstream analysis using Perseus (123). Proteins identified in less than 60% of samples were filtered from the dataset and missing values were imputed. Principal component analysis (PCA), histograms, profile plots and volcano plots were generated using default parameters in Perseus and plotted in R using ggplot2 (https://ggplot2.tidyverse.org/). Significance of fold-changes between experimental conditions were assessed using significance analysis of microarrays (SAM) analysis (S0 = 0.1) with a permutation-based false discovery rate (FDR) of 0.05 to calculate FDR-adjusted p-value (q-value) (124). The Database for Annotation, Visualisation and Integrated Discovery (DAVID) v2025_2 (125) was used to perform GO analysis on genes with significantly altered abundance against a background of all human genes identified in the experiment.

### Quantitative temporal transcriptomics

1.75×10^6^ day 3 i3Neurones were seeded in PLO-treated 6-well plates (1 well per condition per timepoint per replicate) and matured to day 14 before being synchronously infected (MOI 5) with WT or ΔpUL56 HSV1, or mock-infected. Inoculum was removed after 1 h and cells were washed thrice with PBS before being overlaid with conditioned media. At 3, 6, 12, 18 and 30 hpi, medium was removed and cells were washed once with PBS, which was subsequently aspirated and plates frozen at -70°C. RNA was extracted using a PureLink RNA Mini extraction kit (Invitrogen), quantified using a Qubit RNA High Sensitivity (HS) Assay Kit (Invitrogen), and prepared for Oxford Nanopore Technology (ONT) sequencing using the cDNA-PCR Sequencing V14 Barcoding kit (Oxford Nanopore) using 500 ng of RNA per sample and a randomised allocation of barcodes. The concentration of the resultant barcoded cDNA was measured using a Qubit dsDNA HS Assay kit (Invitrogen) and its integrity assessed using an Agilent Bioanalyser 2100 expert High Sensitivity DNA kit (Agilent) before being pooled 1:1 for all labelled samples to a total of 50 fmol per library. Rapid adaptors were added and the libraries were sequenced using four PromethION flow cells (Oxford Nanopore), with each library split across two flow cells. Sequencing ran for three days and base-calling (Super-accurate basecalling v4.3.0, 400 bases per second) was performed in parallel using the Guppy recurrent neural network algorithm (version 7.4.14+d47971cbb) provided by ONT. Run data was then summarised by PycoQC (126), sequence data was combined and the data from across the technical replicates was merged. A combined reference cDNA transcriptome file was created containing human (Ensembl v113) and HSV-1 cDNAs. Minimap 2 (version 2.24-r1122) was used to align ONT reads to the combined reference cDNA transcriptome and aligned reads. Salmon version 1.6.0 (127, 128) was used to assign reads to transcripts or gene isoforms and produce a read count matrix. Differential gene expression was performed in R (Software S2) using DESeq2 from Bioconductor (129). Raw gene-level counts were imported, and genes with low counts across all samples (median count ≤ 3) filtered out prior to normalisation. The DESeq2 median-of-ratios method was employed to calculate size factors, correcting for variations in sequencing depth and RNA composition bias. Variance estimation and dispersion fitting were conducted utilising an empirical Bayes shrinkage procedure. To stabilise variance across the dynamic range, counts were transformed using a variance stabilising transformation before t-Distributed Stochastic Neighbour Embedding (t-SNE) analysis. Differential expression analysis was conducted using Wald tests, with resulting p-values adjusted for multiple testing via the Benjamini–Hochberg FDR procedure. Profile plots, volcano plots and abundance plots were generated using ggplot2 and GO analysis was performed using (DAVID) v2025_2 (125) with a background of all human genes identified in the experiment.

### Plasma membrane proteomics

1×10^7^ day 3 i3Neurones were seeded in PLO-treated 10 cm dishes and matured to day 14 before being synchronously infected (MOI 5) with WT or ΔpUL56 HSV1, or mock-infected. Inoculum was removed after 1 h and cells were washed thrice with PBS before being overlaid with conditioned media. At 24 hpi, cells were washed thrice with ice-cold PBS pH 7.4 then overlain with 5 mL ice-cold PBS supplemented with 1 mM sodium periodate (SigmaAldrich), 100 μM aminooxy-biotin (Biotium) and 10 mM aniline (Sigma-Aldrich). Samples were incubated for 30 min at 4°C protected from light before biotinylation was quenched by addition of an equal volume of 2 mM glycerol. Cells were gently washed thrice with ice-cold PBS and then scraped into 1 mL of ice-cold lysis buffer (10 mM Tris pH 7.4, 1% v/v Triton X100, 150 mM NaCl, 5 mM EDTA and 1% v/v EDTA-free protease inhibitor cocktail) and transferred to low-bind microcentrifuge tubes (Corning). Cells were lysed for 1 h and lysates clarified by centrifugation (20,000 × g, 10 min, 4°C), snap-frozen in liquid nitrogen and stored at -70°C until required. Once all samples were ready for enrichment, lysates were thawed, their protein concentrations measured via bicinchoninic acid assay (Pierce), and protein concentrations were normalised via addition of lysis buffer. Neutravidin beads were washed with lysis buffer before being incubated with samples for 3 h at 4°C. Beads with enriched samples were washed in SnapCap filter columns (Pierce) on a vacuum manifold 20 times using lysis buffer, 20 times using PBS with 0.5% SDS, and then 10 times with urea buffer comprising 6 M urea and 50 mM triethylammonium bicarbonate (TEAB) pH 8.5 (ThermoFisher). Columns were removed, capped, and beads incubated with a reduction/alkylation mixture comprising 10 mM Tris(2-carboxyethyl)phosphine (ThermoFisher) and 20 mM iodoacetamide in urea buffer for 30 min on a shaking platform protected from light. The beads were then washed on the vacuum manifold a further 10 times with urea buffer, then 5 times with 50 mM TEAB. Beads were transferred to a microcentrifuge tube, supernatant was removed and beads were resuspended in 50 μL 50 mM TEAB containing 0.5 μg sequencing-grade trypsin (Promega). Proteins were digested overnight at 37°C in an incubator shaking at 1000 rpm. The next day, beads were pelleted (500 × g, 5 min) and the supernatant reserved before the beads were washed with 40 μL 50 mM TEAB and pelleted again. The wash supernatant was combined with the reserved supernatant, and samples dried in a vacuum centrifuge.

Digested peptides were subjected to LC/MS2 analysis using a Thermo Q Exactive Plus mass spectrometer (ThermoFisher Scientific) equipped with an EASYspray source and coupled to an RSLC3000 nano UPLC (Thermo Fischer Scientific). Peptides were fractionated using a 50 cm C18 PepMap EASYspray column maintained at 40°C with a solvent flow rate of 300 nL/min. A gradient was formed using solvent A (0.1% formic acid) and solvent B (80% v/v acetonitrile, 0.1% formic acid) rising from 3% to 40% solvent B by 90 min followed by a 5 min wash at 95% solvent B. MS spectra were acquired at 35,000 resolution between m/z 350 and 1430 with MSMS spectra acquired in a DIA fashion. Fragmentation spectra were acquired at 17,500 fwhm following HCD activation with a 30 m/z isolation window, 1.0×10^6^ automatic gain control (AGC) target and a loop count of 18. DIA-NN version 1.9.2 (130) was used to analyse the resultant spectra, which were searched against an *in silico* library generated using default parameters from a curated HSV-1 strain KOS proteome and the human proteome (UP000005640, accessed on 06.12.2024). Optimal analysis parameters (mass accuracy and MS1 accuracy) were calculated from a short initial processing of three random samples and then applied to all samples with match between runs, as recommended for proteomic spectra captured on a Q Exactive Plus mass spectrometer. Protein abundances were exported as MaxLFQ scores (131). DIA-NN data were preprocessed in R by removing common contaminant proteins and applying a log2 transformation (Software S3). PMP data was further filtered (Software S4) to only include proteins annotated within biomaRt (132) with GO cellular compartment terms plasma membrane (GO:0005886), cell surface (GO:0009986), or extracellular region (GO:0005576) before analysis using Perseus (123) as above.

### Calcium ion flux imaging

2.5×10^4^ day 3 i3Neurones were seeded per well in a PLO-coated 96-well plate, with three technical triplicate wells for each infection condition and timepoint. After 48 h, 100 μL of medium was removed and 100 μL of Neuroburst Orange Lentivirus (Sartorius), diluted 1:100 in CN medium, was added. At 24 hpi the inoculum was replaced with fresh CN medium, and subsequently half-medium changes were performed thrice weekly. At day 20 the plate was transferred to an Incucyte SX5 for daily imaging with movie mode (2 min/well, 3 images/s, 4× lens). Movie analysis was performed using the Neuronal Activity Analysis software with the following parameters: object size 100 μm; min cell width 4 μm; sensitivity -2; edge split sensitivity -15; min burst intensity 0.05. At day 40, Neuroburst-transduced i3Neurones were synchronously infected (MOI 5) with WT, ΔpUL56 or pUL56 AAXA123 HSV-1, or mock-infected. After 1 h cells were washed thrice with PBS before being overlaid with conditioned media. To prevent bleaching of the calcium biosensor, wells were scanned only once during the 48 h infection time course, either at 12, 24 or 48 hpi. 1 μM tetrodotoxin (TTX) was added at 50 hpi to wells initially scanned at 12 hpi, which were rescanned an hour later. Wells at 24 hpi were fixed and stained for ICP4 to measure percentage infection as described in (27).

### Multielectrode array (MEA) analysis

Wells of a 24-well MEA cytoview plate (Axion Biosystems) were coated with PLO before being seeded with 4×10^5^ day 3 i^3^Neurones. Neurones were matured with half-media changes thrice weekly. From day 14, electrical activity was recorded once every weekday on a Maestro Edge system (Axion Biosystems) with the control software Navigator (Axion Biosystems). The threshold for spike detection was 6 times the standard deviation of the background signal. Before recordings, plates were loaded into the instrument and equilibrated for 15 min at 37°C, 5% CO_2_. Data were processed with Navigator and activity metrics calculated with Neural Metrics tool (Axion Biosystems). Statistical analysis was performed using Prism 7 (GraphPad), comparing conditions via one-way ANOVA and Sidak’s multiple comparisons test.

### Data, Materials, and Software Availability

The mass spectrometry proteomics data produced in this study have been deposited to the ProteomeXchange Consortium via the PRIDE repository (133) (https://www.ebi.ac.uk/pride/) with the dataset identifiers PXD076419 (for the whole-cell proteomics) and PXD076394 (for the plasma membrane proteomics). The transcriptomics data have been deposited in ENA (https://www.ebi.ac.uk/ena/browser/home) under the accession number XXXX. Materials will be shared upon request under a Materials Transfer Agreement. All other supporting data, code and protocols have been provided within the article or through supplementary data files.

## Supporting information

Software S3

Software S4

Software S1

Software S2

## Acknowledgments

We thank Caitlin Gillespie for technical assistance and Michael Coleman for helpful discussions. D.A.N. was supported by a Department of Pathology studentship funded by the Gwynaeth Pretty Fund. This work was supported by a Sir Henry Dale Fellowship (220620/Z/20/Z) from the Wellcome Trust and the Royal Society to V.L., a Wellcome Discovery Award (309425/Z/24/Z) and MRC project grant (MR/X000516/1) to M.P.W., a Wellcome Trust Senior Research Fellowship (219447/Z/19/Z) to J.E.D., and a pump-priming award from the Department of Pathology Bioinformatics and Genomics Trust Fund to S.C.G.. The funders had no role in study design, data collection and analysis, decision to publish or preparation of the manuscript. For the purpose of open access, the authors have applied a Creative Commons Attribution (CC BY) license to any Author Accepted Manuscript version arising from this submission.

## References

1. M. Harfouche, et al., Estimated global and regional incidence and prevalence of herpes simplex virus infections and genital ulcer disease in 2020: mathematical modelling analyses. Sex Transm Infect 101, 214–223 (2025).

2. M. P. Nicoll, J. T. Proença, S. Efstathiou, The molecular basis of herpes simplex virus latency. FEMS Microbiol Rev 36, 684–705 (2012).

3. R. J. Whitley, Herpes simplex encephalitis: adolescents and adults. Antiviral Res 71, 141–148 (2006).

4. M. J. Bradshaw, A. Venkatesan, Herpes Simplex Virus-1 Encephalitis in Adults: Pathophysiology, Diagnosis, and Management. Neurotherapeutics 13, 493–508 (2016).

5. A. Venkatesan, B. D. Michael, J. C. Probasco, R. G. Geocadin, T. Solomon, Acute encephalitis in immunocompetent adults. Lancet 393, 702–716 (2019).

6. L. K. Jørgensen, L. S. Dalgaard, L.J. Østergaard, M. Nørgaard, T. H. Mogensen, Incidence and mortality of herpes simplex encephalitis in Denmark: A nationwide registry-based cohort study. J Infect 74, 42–49 (2017).

7. L. S. Duerlund, et al., Herpes simplex virus type 1 encephalitis: A prospective population-based cohort study. Clin Infect Dis ciag136 (2026). 10.1093/cid/ciag136.

8. R. J. Whitley, et al., Adenine arabinoside therapy of biopsy-proved herpes simplex encephalitis. National Institute of Allergy and Infectious Diseases collaborative antiviral study. N Engl J Med 297, 289–294 (1977).

9. N. D. Rocha, S. K. de Moura, G. A. B. da Silva, R. Mattiello, D. K. Sato, Neurological sequelae after encephalitis associated with herpes simplex virus in children: systematic review and meta-analysis. BMC Infect Dis 23, 55 (2023).

10. J. P. Stahl, A. Mailles, T. De Broucker, Steering Committee and Investigators Group, Herpes simplex encephalitis and management of acyclovir in encephalitis patients in France. Epidemiol Infect 140, 372–381 (2012).

11. P. Jain, et al., Epidemiology and etiology of acute encephalitis syndrome in North India. Jpn J Infect Dis 67, 197–203 (2014).

12. K. Lopatko Lindman, et al., Herpesvirus infections, antiviral treatment, and the risk of dementia-a registry-based cohort study in Sweden. Alzheimers Dement (N Y) 7, e12119 (2021).

13. K. Araya, R. Watson, K. Khanipov, G. Golovko, G. Taglialatela, Increased risk of dementia associated with herpes simplex virus infections: Evidence from a retrospective cohort study using U.S. electronic health records. J Alzheimers Dis 104, 393–402 (2025).

14. Y. Liu, C. Johnston, N. Jarousse, S. P. Fletcher, S. Iqbal, Association between herpes simplex virus type 1 and the risk of Alzheimer’s disease: a retrospective case-control study. BMJ Open 15, e093946 (2025).

15. R. F. Itzhaki, Corroboration of a Major Role for Herpes Simplex Virus Type 1 in Alzheimer’s Disease. Front Aging Neurosci 10, 324 (2018).

16. D. M. Cairns, et al., A 3D human brain-like tissue model of herpes-induced Alzheimer’s disease. Sci Adv 6, eaay8828 (2020).

17. T. K. Soh, et al., Temporal Proteomic Analysis of Herpes Simplex Virus 1 Infection Reveals Cell-Surface Remodeling via pUL56-Mediated GOPC Degradation. Cell Rep 33, 108235 (2020).

18. T. Hennig, L. Djakovic, L. Dölken, A. W. Whisnant, A Review of the Multipronged Attack of Herpes Simplex Virus 1 on the Host Transcriptional Machinery. Viruses 13, 1836 (2021).

19. K. M. Scherer, et al., A fluorescent reporter system enables spatiotemporal analysis of host cell modification during herpes simplex virus-1 replication. Journal of Biological Chemistry 296 (2021).

20. P. N. Canova, A. J. Charron, D. A. Leib, Models of Herpes Simplex Virus Latency. Viruses 16, 747 (2024).

21. H. Fu, D. Pan, Mechanisms of HSV gene regulation during latency and reactivation. Virology 602, 110324 (2025).

22. M. T. Hussain, B. A. Stanfield, D. I. Bernstein, Small Animal Models to Study Herpes Simplex Virus Infections. Viruses 16, 1037 (2024).

23. A. Rybak-Wolf, et al., Modelling viral encephalitis caused by herpes simplex virus 1 infection in cerebral organoids. Nat Microbiol 8, 1252–1266 (2023).

24. C. Wang, et al., Scalable Production of iPSC-Derived Human Neurons to Identify Tau-Lowering Compounds by High-Content Screening. Stem Cell Reports 9, 1221–1233 (2017).

25. M. S. Fernandopulle, et al., Transcription Factor-Mediated Differentiation of Human iPSCs into Neurons. Curr Protoc Cell Biol 79, e51 (2018).

26. A. H. M. Ng, et al., A comprehensive library of human transcription factors for cell fate engineering. Nat Biotechnol 39, 510–519 (2021).

27. D. A. Nash, et al., Optimization of lytic herpes simplex virus infection in human induced pluripotent stem cell-derived cortical neurones. Journal of General Virology 107, 002237 (2026).

28. H. S. Oh, et al., Validation of human sensory neurons derived from inducible pluripotent stem cells as a model for latent infection and reactivation by herpes simplex virus 1. mBio 16, e0187125 (2025).

29. Y. Deng, et al., Neuronal miR-9 promotes HSV-1 epigenetic silencing and latency by repressing Oct-1 and Onecut family genes. Nat Commun 15, 1991 (2024).

30. A. S. Nicholson, et al., Plasma membrane remodeling in GM2 gangliosidoses drives synaptic dysfunction. PLoS Biol 23, e3003265 (2025).

31. H. G. Barrow, et al., Human iPSC Models of Ganglioside Deficiency Reveal a Sialylated Lipid Requirement for Plasma-Membrane Organization and Neuronal Activity. [Preprint] (2026). Available at: http://biorxiv.org/lookup/doi/10.64898/2026.03.18.712603 [Accessed 1 April 2026].

32. A. Fletcher-Etherington, M. P. Weekes, Quantitative Temporal Viromics. Annu Rev Virol 8, 159–181 (2021).

33. K. Nightingale, et al., High-Definition Analysis of Host Protein Stability during Human Cytomegalovirus Infection Reveals Antiviral Factors and Viral Evasion Mechanisms. Cell Host Microbe 24, 447-460.e11 (2018).

34. S. Ozon, A. Maucuer, A. Sobel, The stathmin family -- molecular and biological characterization of novel mammalian proteins expressed in the nervous system. Eur J Biochem 248, 794–806 (1997).

35. I. Bièche, et al., Expression of stathmin family genes in human tissues: non-neural-restricted expression for SCLIP. Genomics 81, 400–410 (2003).

36. Z. J. Gieroba, B.-S. Zhu, W. W. Blessing, S. L. Wesselingh, Herpes Simplex Virus induces Fos expression in rat brainstem neurons. Brain Research 675, 329–332 (1995).

37. B. Hu, et al., Cellular responses to HSV-1 infection are linked to specific types of alterations in the host transcriptome. Sci Rep 6, 28075 (2016).

38. R. Hagglund, C. Van Sant, P. Lopez, B. Roizman, Herpes simplex virus 1-infected cell protein 0 contains two E3 ubiquitin ligase sites specific for different E2 ubiquitin-conjugating enzymes. Proc Natl Acad Sci U S A 99, 631–636 (2002).

39. C. Boutell, S. Sadis, R. D. Everett, Herpes simplex virus type 1 immediate-early protein ICP0 and is isolated RING finger domain act as ubiquitin E3 ligases in vitro. J Virol 76, 841–850 (2002).

40. K. M. Eidson, W. E. Hobbs, B. J. Manning, P. Carlson, N. A. DeLuca, Expression of herpes simplex virus ICP0 inhibits the induction of interferon-stimulated genes by viral infection. J Virol 76, 2180–2191 (2002).

41. T. Koshizuka, T. Kobayashi, K. Ishioka, T. Suzutani, Herpesviruses possess conserved proteins for interaction with Nedd4 family ubiquitin E3 ligases. Sci Rep 8, 4447 (2018).

42. A. E. Cuadra, S.-H. Kuo, Y. Kawasaki, D. S. Bredt, D. M. Chetkovich, AMPA Receptor Synaptic Targeting Regulated by Stargazin Interactions with the Golgi-Resident PDZ Protein nPIST. J. Neurosci. 24, 7491–7502 (2004).

43. N. Tongmuang, M. Krishnan, V. Connor, C. Crump, L. E. Jensen, UL56 Is Essential for Herpes Simplex Virus-1 Virulence In Vivo but Is Dispensable for Induction of Host-Protective Immunity. Vaccines (Basel) 12, 837 (2024).

44. T. Kanematsu, et al., Role of the PLC-related, catalytically inactive protein p130 in GABA(A) receptor function. EMBO J 21, 1004–1011 (2002).

45. K. Takenaka, et al., Role of phospholipase C-L2, a novel phospholipase C-like protein that lacks lipase activity, in B-cell receptor signaling. Mol Cell Biol 23, 7329–7338 (2003).

46. E. V. Gazina, et al., “Neonatal” Nav1.2 reduces neuronal excitability and affects seizure susceptibility and behaviour. Hum Mol Genet 24, 1457–1468 (2015).

47. K. M. Stiles, R. S. B. Milne, G. H. Cohen, R. J. Eisenberg, C. Krummenacher, The herpes simplex virus receptor nectin-1 is down-regulated after trans-interaction with glycoprotein D. Virology 373, 98–111 (2008).

48. H. Tanno, T. Yamaguchi, E. Goto, S. Ishido, M. Komada, The Ankrd 13 family of UIM-bearing proteins regulates EGF receptor endocytosis from the plasma membrane. Mol Biol Cell 23, 1343–1353 (2012).

49. Q. Zhang, M. Martin-Caraballo, S. V. Hsia, Modulation of Voltage-Gated Sodium Channel Activity in Human Dorsal Root Ganglion Neurons by Herpesvirus Quiescent Infection. J Virol 94, e01823–19 (2020).

50. N. Storey, D. Latchman, S. Bevan, Selective internalization of sodium channels in rat dorsal root ganglion neurons infected with herpes simplex virus-1. J Cell Biol 158, 1251–1262 (2002).

51. S. G. Oakes, R. W. Petry, R. J. Ziegler, R. S. Pozos, Electrophysiological changes of HSV-1-infected dorsal root ganglia neurons in culture. J Neuropathol Exp Neurol 40, 380–389 (1981).

52. J. Fukuda, T. Kurata, K. Yamaguchi, Specific reduction in Na currents after infection with herpes simplex virus in cultured mammalian nerve cells. Brain Res 268, 367–371 (1983).

53. Q. Zhang, S.-C. Hsia, M. Martin-Caraballo, Regulation of T-type Ca2+ channel expression by interleukin-6 in sensory-like ND7/23 cells post-herpes simplex virus (HSV-1) infection. J Neurochem 151, 238–254 (2019).

54. Q. Zhang, S.-C. Hsia, M. Martin-Caraballo, Regulation of T-type Ca2+ channel expression by herpes simplex virus-1 infection in sensory-like ND7 cells. J Neurovirol 23, 657–670 (2017).

55. R. Piacentini, et al., Herpes Simplex Virus type-1 infection induces synaptic dysfunction in cultured cortical neurons via GSK-3 activation and intraneuronal amyloid-β protein accumulation. Sci Rep 5, 15444 (2015).

56. F. Acuña-Hinrichsen, et al., Herpes Simplex Virus Type 1 Neuronal Infection Triggers the Disassembly of Key Structural Components of Dendritic Spines. Front. Cell. Neurosci. 15 (2021).

57. P. Bergström, et al., Herpes Simplex Virus 1 and 2 Infections during Differentiation of Human Cortical Neurons. Viruses 13, 2072 (2021).

58. K. M. McCarthy, D. W. Tank, L. W. Enquist, Pseudorabies Virus Infection Alters Neuronal Activity and Connectivity In Vitro. PLoS Pathog 5, e1000640 (2009).

59. R. Volmer, C. Monnet, D. Gonzalez-Dunia, Borna Disease Virus Blocks Potentiation of Presynaptic Activity through Inhibition of Protein Kinase C Signaling. PLOS Pathogens 2, e19 (2006).

60. J. Gaburro, et al., Zika virus-induced hyper excitation precedes death of mouse primary neuron. Virol J 15, 79 (2018).

61. S. Ghassemi, et al., Lentiviral Expression of Rabies Virus Glycoprotein in the Rat Hippocampus Strengthens Synaptic Plasticity. Cell Mol Neurobiol 42, 1429–1440 (2022).

62. P. R. Kramer, M. Umorin, R. Hornung, P. R. Kinchington, Neurexin 3α in the central amygdala has a role orofacial varicella zoster pain. Neuroscience 496, 16–26 (2022).

63. M. V. Green, J. D. Raybuck, X. Zhang, M. M. Wu, S. A. Thayer, Scaling Synapses in the Presence of HIV. Neurochem Res 44, 234–246 (2019).

64. J. Jędrzejewska-Szmek, D. B. Dorman, K. T. Blackwell, Making Time and Space for Calcium Control of Neuron Activity. Curr Opin Neurobiol 83, 102804 (2023).

65. Z. Li, I. P. Michael, D. Zhou, A. Nagy, J. M. Rini, Simple piggyBac transposon-based mammalian cell expression system for inducible protein production. Proc Natl Acad Sci U S A 110, 5004–5009 (2013).

66. R. De Pace, et al., Messenger RNA transport on lysosomal vesicles maintains axonal mitochondrial homeostasis and prevents axonal degeneration. Nat Neurosci 27, 1087–1102 (2024).

67. E. J. C. Thornburg-Suresh, D. W. Summers, Microtubules, membranes, and movement: New roles for Stathmin-2 in axon integrity. J Neurosci Res 102, e25382 (2024).

68. J. E. Shin, S. Geisler, A. DiAntonio, Dynamic regulation of SCG10 in regenerating axons after injury. Exp Neurol 252, 1–11 (2014).

69. Y. Okada, H. Yamazaki, Y. Sekine-Aizawa, N. Hirokawa, The neuron-specific kinesin superfamily protein KIF1A is a unique monomeric motor for anterograde axonal transport of synaptic vesicle precursors. Cell 81, 769–780 (1995).

70. M. P. Coleman, A. Höke, Programmed axon degeneration: from mouse to mechanism to medicine. Nat Rev Neurosci 21, 183–196 (2020).

71. Z. Melamed, et al., Premature polyadenylation-mediated loss of stathmin-2 is a hallmark of TDP-43-dependent neurodegeneration. Nat Neurosci 22, 180–190 (2019).

72. K. L. Krus, et al., Loss of Stathmin-2, a hallmark of TDP-43-associated ALS, causes motor neuropathy. Cell Rep 39, 111001 (2022).

73. J. López-Erauskin, et al., Stathmin-2 loss leads to neurofilament-dependent axonal collapse driving motor and sensory denervation. Nat Neurosci 27, 34–47 (2024).

74. D. Freisem, et al., Herpes simplex virus infection promotes ALS pathology through ICP0-mediated PML body disruption. [Preprint] (2026). Available at: http://biorxiv.org/lookup/doi/10.64898/2026.03.27.714707 [Accessed 9 April 2026].

75. T. Koshizuka, Y. Kawaguchi, Y. Nishiyama, Herpes simplex virus type 2 membrane protein UL56 associates with the kinesin motor protein KIF1A. J Gen Virol 86, 527–533 (2005).

76. J. Scherer, et al., A kinesin-3 recruitment complex facilitates axonal sorting of enveloped alpha herpesvirus capsids. PLoS Pathog 16, e1007985 (2020).

77. T. Kramer, et al., Kinesin-3 mediates axonal sorting and directional transport of alphaherpesvirus particles in neurons. Cell Host Microbe 12, 806–814 (2012).

78. J. Kumar, et al., The Caenorhabditis elegans Kinesin-3 motor UNC-104/KIF1A is degraded upon loss of specific binding to cargo. PLoS Genet 6, e1001200 (2010).

79. J. Cisneros Solis, T. L. Blasius, K. J. Verhey, The Kinesin-3 KIF1A Functions as a Diligent Worker during Axonal Transport. [Preprint] (2025). Available at: http://biorxiv.org/lookup/doi/10.1101/2025.09.11.675527 [Accessed 2 April 2026].

80. A. Negatsch, T. C. Mettenleiter, W. Fuchs, Herpes simplex virus type 1 strain KOS carries a defective US9 and a mutated US8A gene. Journal of General Virology 92, 167–172 (2011).

81. P. W. Howard, T. L. Howard, D. C. Johnson, Herpes simplex virus membrane proteins gE/gI and US9 act cooperatively to promote transport of capsids and glycoproteins from neuron cell bodies into initial axon segments. J Virol 87, 403–414 (2013).

82. N. Hensel, et al., The Proteome and Secretome of Cortical Brain Cells Infected With Herpes Simplex Virus. Front Neurol 11, 844 (2020).

83. H. T. W. Blest, et al., HSV-1 employs UL56 to antagonize expression and function of cGAMP channels. Cell Reports 43 (2024).

84. M. X. van Bemmelen, et al., Cardiac voltage-gated sodium channel Nav1.5 is regulated by Nedd4-2 mediated ubiquitination. Circ Res 95, 284–291 (2004).

85. C. J. Laedermann, et al., Dysregulation of voltage-gated sodium channels by ubiquitin ligase NEDD4-2 in neuropathic pain. J Clin Invest 123, 3002–3013 (2013).

86. A. García-Caballero, et al., The deubiquitinating enzyme USP5 modulates neuropathic and inflammatory pain by enhancing Cav3.2 channel activity. Neuron 83, 1144–1158 (2014).

87. K. M. Wright, et al., The C-Terminal of NaV1.7 Is Ubiquitinated by NEDD4L. ACS Bio Med Chem Au 3, 516– 527 (2023).

88. S. Tyagi, et al., Targeted ubiquitination of Na V 1.8 reduces sensory neuronal excitability. bioRxiv 2025.02.04.636451 (2025). 10.1101/2025.02.04.636451.

89. A. B. Fotia, et al., Regulation of neuronal voltage-gated sodium channels by the ubiquitin-protein ligases Nedd4 and Nedd4-2. J Biol Chem 279, 28930–28935 (2004).

90. L. Qiu, et al., HSV-1 UL56 protein recruits cellular NEDD4-family ubiquitin ligases to suppress CD1d expression and NKT cell function. J Virol 99, e0214024 (2025).

91. T. Koshizuka, et al., Identification and characterization of the UL56 gene product of herpes simplex virus type 2. J Virol 76, 6718–6728 (2002).

92. Y. Ushijima, T. Koshizuka, F. Goshima, H. Kimura, Y. Nishiyama, Herpes Simplex Virus Type 2 UL56 Interacts with the Ubiquitin Ligase Nedd4 and Increases Its Ubiquitination. Journal of Virology 82, 5220–5233 (2008).

93. H. Jiang, S. N. Thomas, Z. Chen, C. Y. Chiang, P. A. Cole, Comparative analysis of the catalytic regulation of NEDD4-1 and WWP2 ubiquitin ligases. J Biol Chem 294, 17421–17436 (2019).

94. M. Tao, et al., ITCH directly K63-ubiquitinates the NOD2 binding protein, RIP2, to influence inflammatory signaling pathways. Curr Biol 19, 1255–1263 (2009).

95. M. Wainberg, et al., The viral hypothesis: how herpesviruses may contribute to Alzheimer’s disease. Mol Psychiatry 26, 5476–5480 (2021).

96. G. A. Jamieson, N. J. Maitland, G. K. Wilcock, J. Craske, R. F. Itzhaki, Latent herpes simplex virus type 1 in normal and Alzheimer’s disease brains. J Med Virol 33, 224–227 (1991).

97. Y. Dai, et al., TMEFF1 is a neuron-specific restriction factor for herpes simplex virus. Nature 632, 383–389 (2024).

98. L. S. Reinert, et al., Sensing of HSV-1 by the cGAS-STING pathway in microglia orchestrates antiviral defence in the CNS. Nat Commun 7, 13348 (2016).

99. S.-Y. Zhang, J.-L. Casanova, Genetic defects of brain immunity in childhood herpes simplex encephalitis. Nature 635, 563–573 (2024).

100. T. J. Baumgartner, et al., Voltage-Gated Na+ Channels in Alzheimer’s Disease: Physiological Roles and Therapeutic Potential. Life (Basel) 13, 1655 (2023).

101. J. Wang, S.-W. Ou, Y.-J. Wang, Distribution and function of voltage-gated sodium channels in the nervous system. Channels (Austin) 11, 534–554 (2017).

102. R. Zhao, et al., Activity disruption causes degeneration of entorhinal neurons in a mouse model of Alzheimer’s circuit dysfunction. eLife 11, e83813 (2022).

103. S. Meftah, J. Gan, Alzheimer’s disease as a synaptopathy: Evidence for dysfunction of synapses during disease progression. Front Synaptic Neurosci 15, 1129036 (2023).

104. A. J. Dutton, et al., Asymptomatic neonatal herpes simplex virus infection in mice leads to persistent CNS infection and long-term cognitive impairment. PLoS Pathog 21, e1012935 (2025).

105. L. Letenneur, et al., Seropositivity to herpes simplex virus antibodies and risk of Alzheimer’s disease: a population-based cohort study. PLoS One 3, e3637 (2008).

106. E. Vestin, et al., Herpes Simplex Viral Infection Doubles the Risk of Dementia in a Contemporary Cohort of Older Adults: A Prospective Study. J Alzheimers Dis 97, 1841–1850 (2024).

107. E. Shin, et al., The associations of herpes simplex virus and varicella zoster virus infection with dementia: a nationwide retrospective cohort study. Alzheimers Res Ther 16, 57 (2024).

108. N.-S. Tzeng, et al., Anti-herpetic Medications and Reduced Risk of Dementia in Patients with Herpes Simplex Virus Infections-a Nationwide, Population-Based Cohort Study in Taiwan. Neurotherapeutics 15, 417– 429 (2018).

109. C. Schnier, et al., Antiherpetic medication and incident dementia: Observational cohort studies in four countries. Eur J Neurol 28, 1840–1848 (2021).

110. K. M. Prasad, et al., Antiherpes virus-specific treatment and cognition in schizophrenia: a test-of-concept randomized double-blind placebo-controlled trial. Schizophr Bull 39, 857–866 (2013).

111. D. P. Devanand, et al., Valacyclovir Treatment of Early Symptomatic Alzheimer Disease: The VALAD Randomized Clinical Trial. JAMA 335, 511–522 (2026).

112. M. Le Hars, C. Joussain, T. Jégu, A. L. Epstein, Non-replicative herpes simplex virus genomic and amplicon vectors for gene therapy - an update. Gene Ther 32, 173–183 (2025).

113. Y. Miyagawa, et al., Herpes simplex viral-vector design for efficient transduction of nonneuronal cells without cytotoxicity. Proc Natl Acad Sci U S A 112, E1632–1641 (2015).

114. S. Cattaneo, et al., Genetic mutations in HSV-1 replication-defective vectors: Implications for their safety in gene therapy applications. Gene Ther 32, 581–593 (2025).

115. T. Russell, B. Bleasdale, M. Hollinshead, G. Elliott, Qualitative Differences in Capsidless L-Particles Released as a By-Product of Bovine Herpesvirus 1 and Herpes Simplex Virus 1 Infections. J Virol 92, e01259–18 (2018).

116. P. Liljeström, S. Lusa, D. Huylebroeck, H. Garoff, In vitro mutagenesis of a full-length cDNA clone of Semliki Forest virus: the small 6,000-molecular-weight membrane protein modulates virus release. J Virol 65, 4107–4113 (1991).

117. K. Rausalu, et al., Properties and use of novel replication-competent vectors based on Semliki Forest virus. Virol J 6, 33 (2009).

118. D. A. McClelland, et al., pH Reduction as a Trigger for Dissociation of Herpes Simplex Virus Type 1 Scaffolds. J Virol 76, 7407–7417 (2002).

119. K. Rausalu, et al., Properties and use of novel replication-competent vectors based on Semliki Forest virus. Virol J 6, 33 (2009).

120. K. J. Livak, T. D. Schmittgen, Analysis of Relative Gene Expression Data Using Real-Time Quantitative PCR and the 2-ΔΔCT Method. Methods 25, 402–408 (2001).

121. S. Tyanova, T. Temu, J. Cox, The MaxQuant computational platform for mass spectrometry-based shotgun proteomics. Nat Protoc 11, 2301–2319 (2016).

122. J. Cox, M. Mann, MaxQuant enables high peptide identification rates, individualized p.p.b.-range mass accuracies and proteome-wide protein quantification. Nat Biotechnol 26, 1367–1372 (2008).

123. S. Tyanova, et al., The Perseus computational platform for comprehensive analysis of (prote)omics data. Nat Methods 13, 731–740 (2016).

124. V. G. Tusher, R. Tibshirani, G. Chu, Significance analysis of microarrays applied to the ionizing radiation response. Proceedings of the National Academy of Sciences 98, 5116–5121 (2001).

125. B. T. Sherman, et al., DAVID: a web server for functional enrichment analysis and functional annotation of gene lists (2021 update). Nucleic Acids Res 50, W216–W221 (2022).

126. A. Leger, T. Leonardi, pycoQC, interactive quality control for Oxford Nanopore Sequencing. Journal of Open Source Software 4, 1236 (2019).

127. H. Li, Minimap2: pairwise alignment for nucleotide sequences. Bioinformatics 34, 3094–3100 (2018).

128. R. Patro, G. Duggal, M. I. Love, R. A. Irizarry, C. Kingsford, Salmon: fast and bias-aware quantification of transcript expression using dual-phase inference. Nat Methods 14, 417–419 (2017).

129. M. I. Love, W. Huber, S. Anders, Moderated estimation of fold change and dispersion for RNA-seq data with DESeq2. Genome Biol 15, 550 (2014).

130. V. Demichev, C. B. Messner, S. I. Vernardis, K. S. Lilley, M. Ralser, DIA-NN: neural networks and interference correction enable deep proteome coverage in high throughput. Nat Methods 17, 41–44 (2020).

131. J. D. O’Connell, J. A. Paulo, J. J. O’Brien, S. P. Gygi, Proteome-Wide Evaluation of Two Common Protein Quantification Methods. J Proteome Res 17, 1934–1942 (2018).

132. S. Durinck, P. T. Spellman, E. Birney, W. Huber, Mapping identifiers for the integration of genomic datasets with the R/Bioconductor package biomaRt. Nat Protoc 4, 1184–1191 (2009).

133. Y. Perez-Riverol, et al., The PRIDE database at 20 years: 2025 update. Nucleic Acids Res 53, D543–D553 (2025).

