## supplementary for "Herpes simplex virus pUL56 abolishes neuronal activity by removing voltage-gated ion channels from the plasma membrane"

**This PDF file includes:**

Figures S1 to S9
Tables S1 to S3
Legends for Movies S1 to S3
Legends for Datasets S1 to S4
Legends for Software S1 to S4
SI References

**Other supporting materials for this manuscript include the following:**

Movies S1 to S3
Datasets S1 to S4
Software S1 to S4

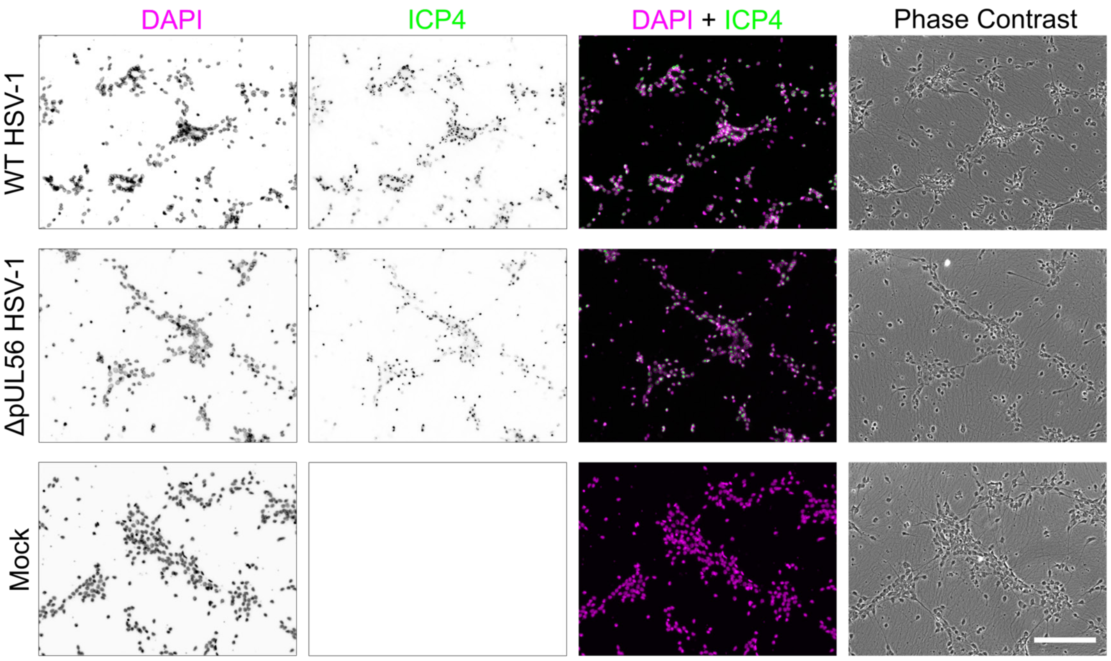


**Fig. S1. i³Neurones synchronously infected with WT and ΔpUL56 HSV-1.**

i³Neurones were infected at MOI 5 with WT and ΔpUL56 HSV-1 concurrently during sample preparation for temporal proteomics. Cells were fixed 18 hpi, immunostained for ICP4 (green) and stained with DAPI (Magenta) to visualise DNA. Co-localisation of ICP4 and DAPI signal confirms near 100% infection. Scale bar = 150um

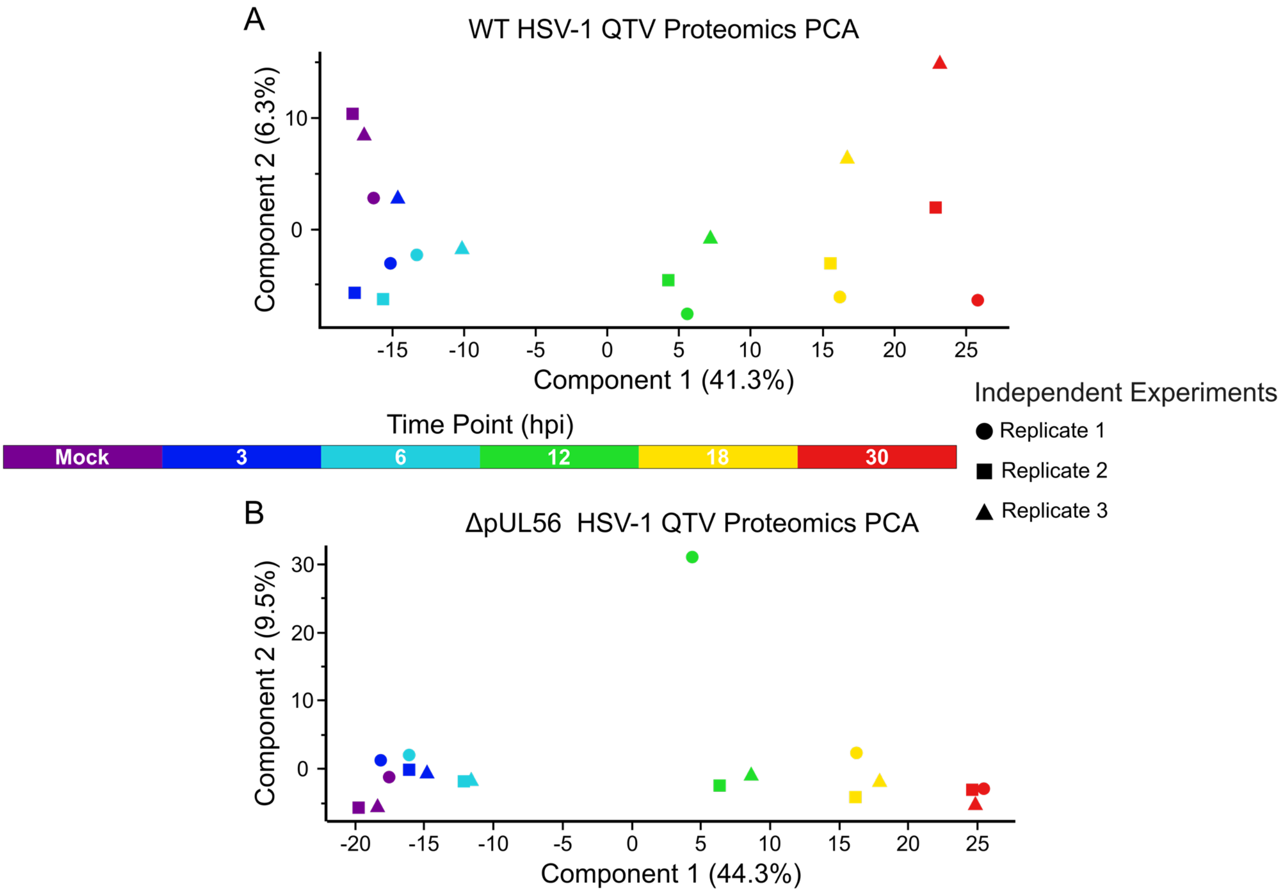


**Fig. S2. Principal component analysis (PCA) of temporal proteomics.**

PCA for (A) WT and (B) ΔpUL56 HSV-1 infection experiments are show, each with their corresponding mock infection. Data points are coloured by time post infection: 3 h (blue), 6 h (cyan), 12 h (green), 18 h (yellow), 30 h (red), and mock infection (purple). Data points cluster by treatment, not biological replicate (squares, circles and triangles). Separation along the horizontal axis for both experiments represents the largest source of variation, corresponding to time post-infection.

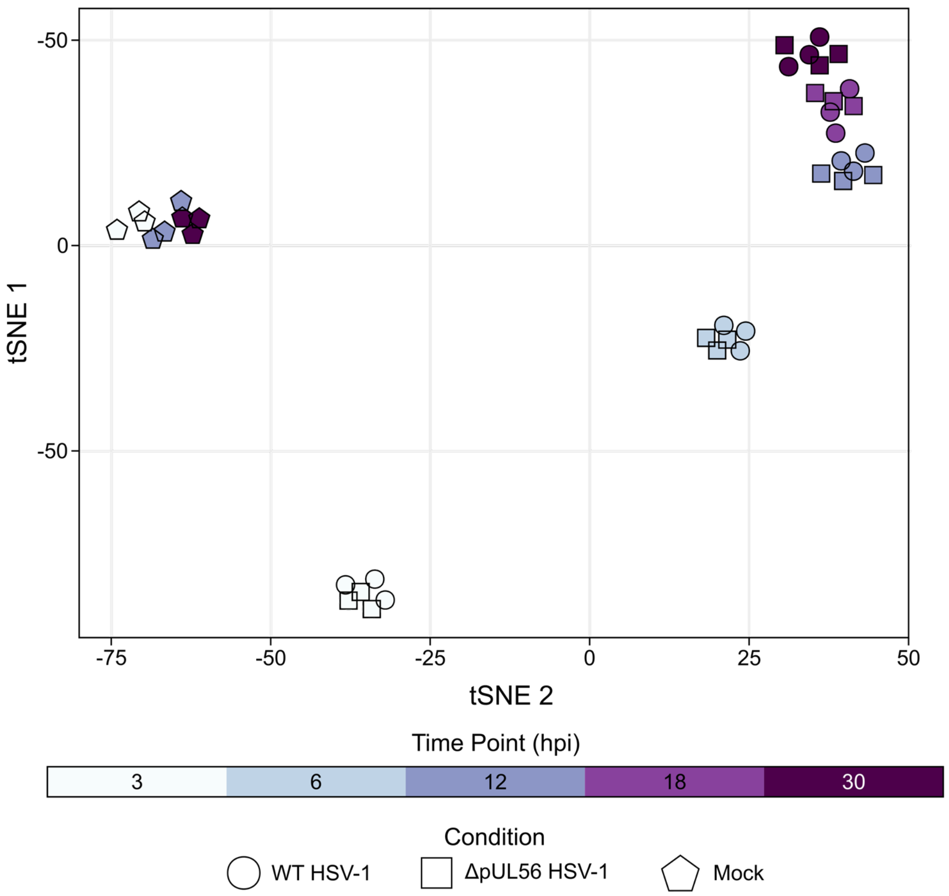


**Fig. S3. T-distributed Stochastic Neighbor Embedding (tSNE) analysis of temporal transcriptomics of WT and ΔpUL56 HSV-1 infected i3Neurones.**

tSNE analysis was performed on variance-stabilised gene counts, coloured by time post (mock)-infection: 3 h (light blue), 6 h (blue), 12 h (slate blue), 18 h (light purple), 30 h (deep purple). Data clusters by treatment and not by biological replicate. At each timepoint WT and ΔpUL56 infection experiments clustering together, consistent with both viruses causing similar changes to the transcriptome of infected i3Neurones.

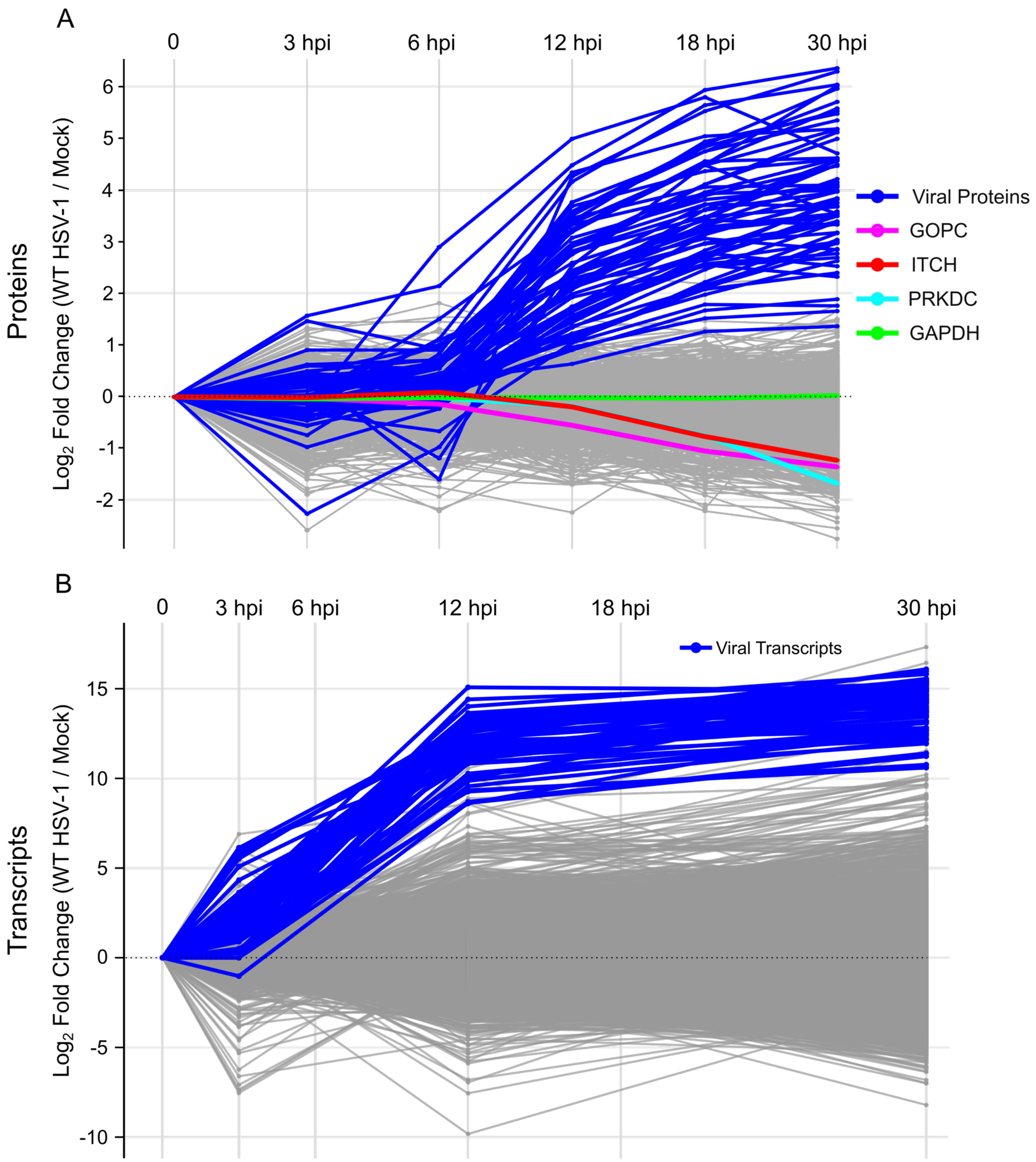


**Fig. S4. Temporal changes in host and virus protein and RNA transcript abundance following infection of i3Neurones with WT HSV-1.**

The mean $\log_2$ fold change in abundance of each (A) protein and (B) transcript is plotted as a function of time post-infection, with mock-infected samples displayed as time 0. Cellular gene products are plotted in grey and viral gene products in blue. (A) $\mathrm{Log}_2$ fold change is calculate relative to the mock-infected sample (taken at 12 hpi). Targets of virus-mediated degradation are highlighted (GOPC, magenta; PRKDC, cyan), as is cFOS that increases in abundance during infection. The abundance of GAPDH (green) does not change during infection. (B) $\mathrm{Log}_2$ fold change is calculated relative to the mock-infected samples taken at the given time points; the 6 hpi and 18 hpi are excluded because mock samples were not taken at those time points. The global decline in mRNA abundance caused by host shutoff is not captured in these plots owing to DESeq2 normalisation that assumes constant total mRNA levels, as discussed in (1).

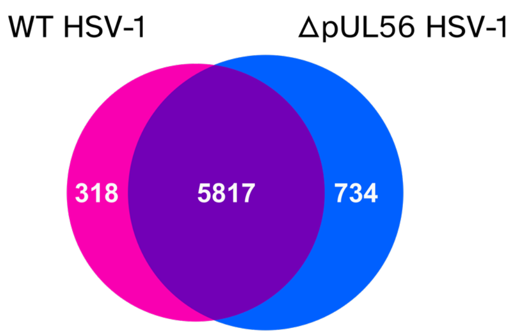


**Fig. S5. Unique protein groups identified in temporal proteomics of i3Neurones infected with WT or ΔpUL56 HSV-1.**

Protein groups can represent more than one protein or isoform if all the identified constituent peptides cannot be unambiguously assigned to a single protein or isoform (3). 5817 protein groups were identified in both experiments, whilst 318 and 734 protein groups were identified only in the WT -1 and ΔpUL56 HSV-1 proteomics analyses, respectively.

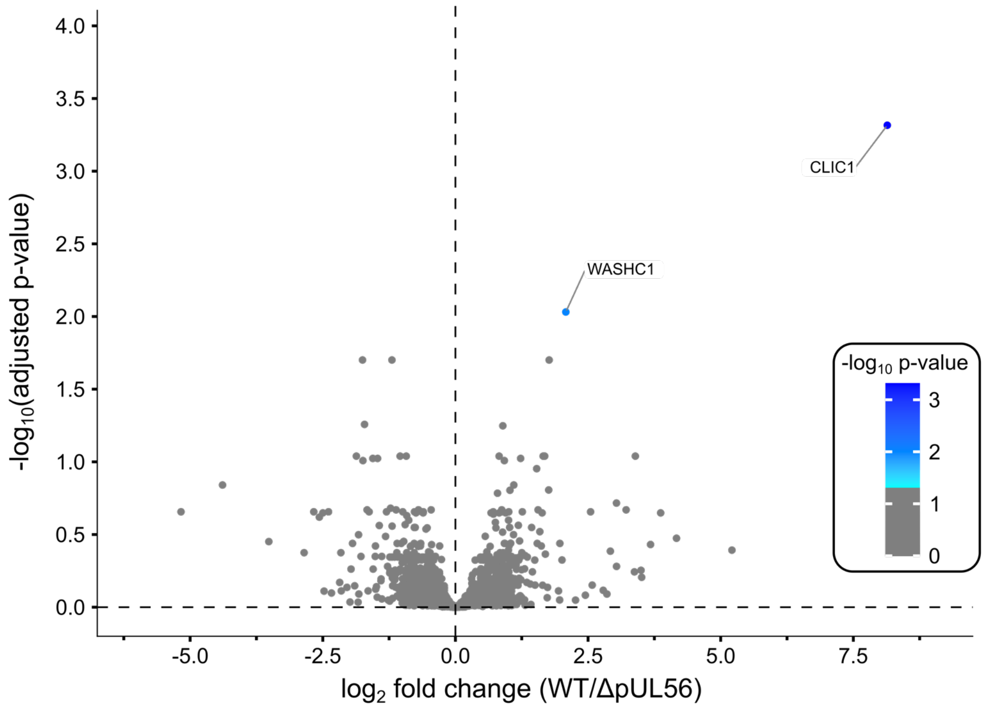


**Fig. S6. Changes in i3Neurone transcript abundance in WT versus ΔpUL56 HSV-1 infected i3Neurones at 30 hpi.**

Horizontal axis shows average $log_2$ fold change and vertical axis shows FDR-adjusted significance for three independent experiments. Significantly altered transcripts ($log_2$ fold change ≥ 2 and p ≤ 0.05) labelled and coloured by significance (blue).

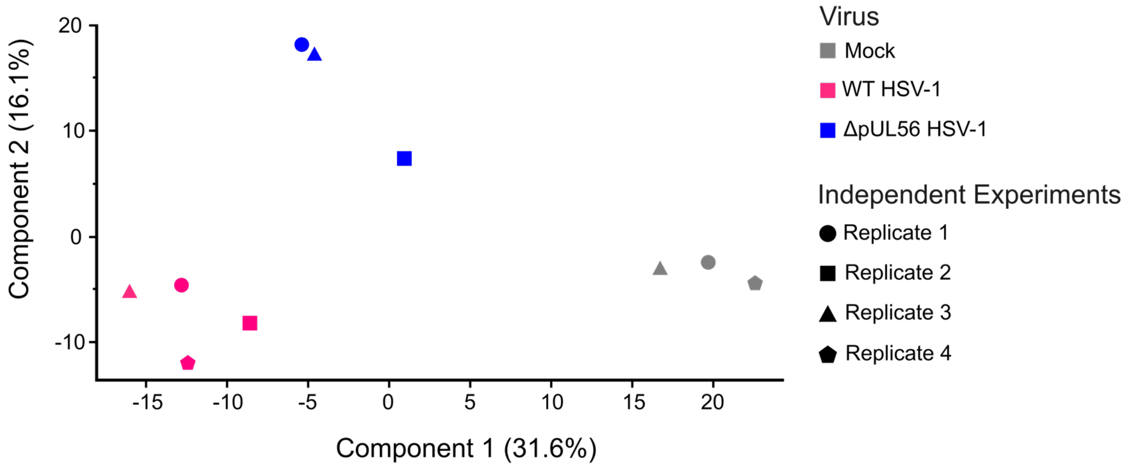


**Fig. S7. PCA of i3Neurone HSV-1 infection PM proteomic analysis.**

PCA was performed on protein abundances that had been $log_2$-transformed, filtered to retain only GO cellular compartment annotations “plasma membrane”, “cell surface”, or “extracellular region”, then processed to impute missing values. Samples cluster by treatment (mock-infection, grey; WT HSV-1 infection, magenta; ΔpUL56 HSV-1 infection, blue) rather than by biological replicate (circles, squares, triangles and pentagons).

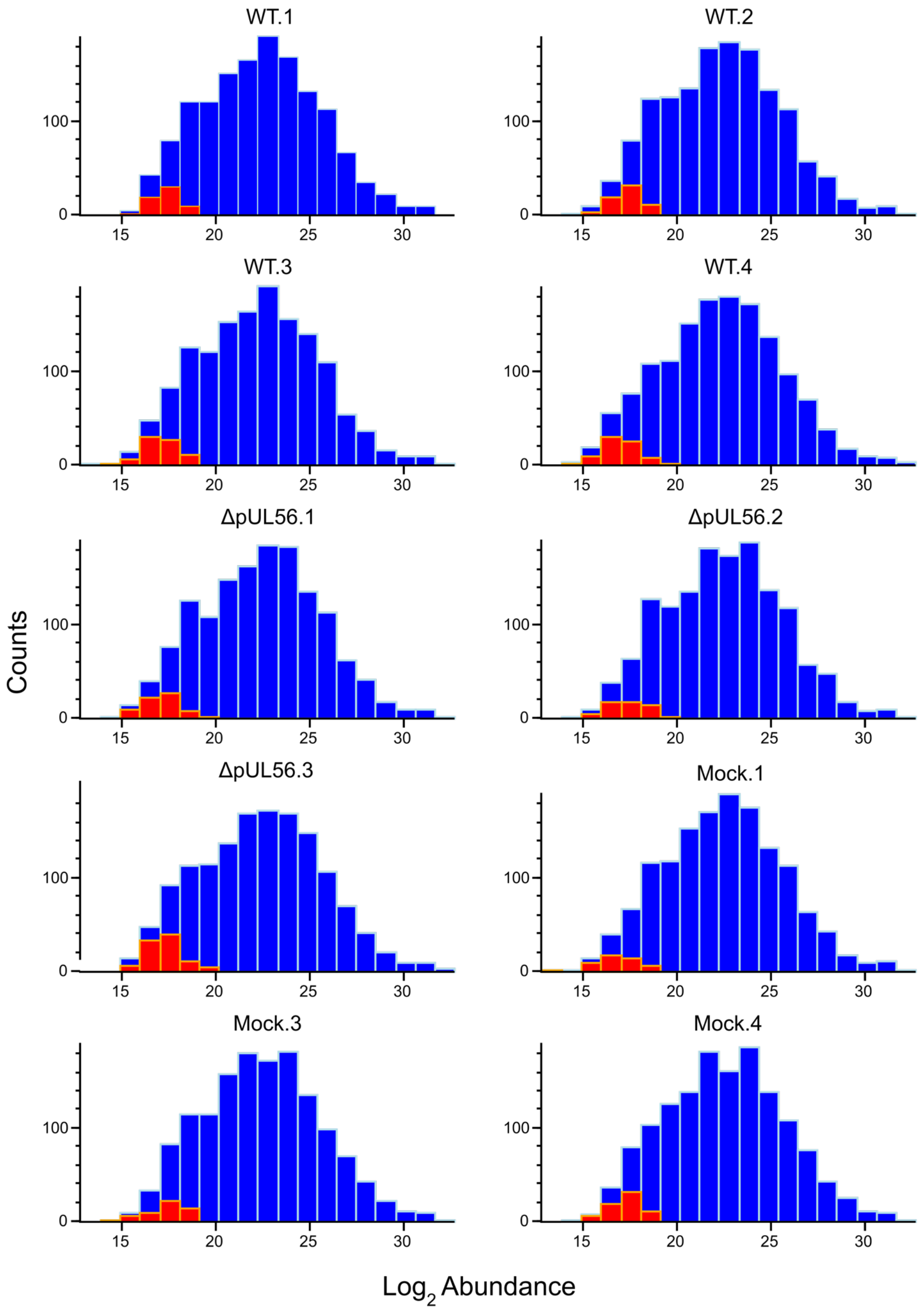


**Fig. S8. PM protein abundances per sample approximately follow a Gaussian distribution after imputation.**

Frequency histogram of observed (blue) and imputed (red) MaxLFQ abundances, $log_2$ transformed, plotted per individual mass spectrometry experiment as shown.

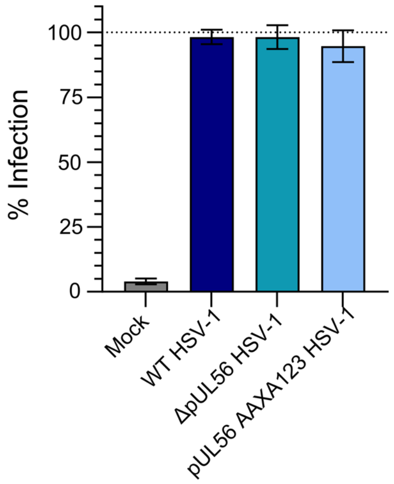


**Fig. S9. i³Neurones were infected with HSV-1 at near-100% efficiency for calcium-flux kinetics assays.**

i3Neurones that had been used for calcium-flux kinetic imaging at 24 hpi were then fixed, immunostained for ICP4 and stained with propidium iodide to visualise HSV-1 infection and DNA, respectively. Co-localisation was quantitated as proportion of propidium iodide positive nuclei with ICP4 staining and is shown as mean ± SD from technical triplicate measurements. For all viruses, approximately 100% of i3Neurones are infected.

**Table S1. Oligonuclotide primers used in this study.**

| Primer | Sequence (5′ to 3′) | Notes |
|---|---|---|
| GOPC KD1 fwd | CACCGACAGAACCGCAGGA GTAACG | Targets GOPC Exon 1 (nt 86–105, antisense), with added BpiI site |
| GOPC KD1 rev | AAACCGTTACTCCTGCGGTT CTGTC | Targets GOPC Exon 1 (nt 86–105, antisense), with added BpiI site |
| GOPC KD2 fwd | CACCGCTGGCGGGGTATCC ATGTTC | Targets GOPC Exon 1 (nt 328–347, sense), with added BpiI site |
| GOPC KD2 rev | AAACGAACATGGATACCCCG CCAGC | Targets GOPC Exon 1 (nt 328–347, sense), with added BpiI site |
| iC10_PB_Internal_FWD | CTGGCGAAAGGGGGATGTG | NEBuilder HiFi assembly of HA-pUL56 piggyBac expression vector |
| iC10_PB_Internal_REV | CACATCCCCCTTTCGCCAG | NEBuilder HiFi assembly of HA-pUL56 piggyBac expression vector |
| Frag1_HA-UL56_FWD | CTCGAGCCACCATGTACCCA TACGATGTTCCAGATTACGCT GCTTCGGAGGCGGCG | NEBuilder HiFi assembly of HA-pUL56 piggyBac expression vector |
| Frag1_HA-UL56_REV | CGTATGGGTACATGGTGGCT CGAGCGGCCGCCGTCGACA TTCTTTGCCAAAATG | NEBuilder HiFi assembly of HA-pUL56 piggyBac expression vector |
| UL56_Frag2_FWD | CTGTGGCGGTGAGAATT CCTTGGACGCGTCTGCAG | NEBuilder HiFi assembly of HA-pUL56 piggyBac expression vector |
| UL56_Frag2_REV | CAAGGAATTCTCACCGCC ACAGGAATACCAG | NEBuilder HiFi assembly of HA-pUL56 piggyBac expression vector |
| Xho1_HA-UL56_FWD | GCGGCCGCTCGAGATGTAC CCATACGATGTTCCAGATTA CGCTGCTTCGGAGGCGGCG | Subcloning pUL56 AAXA123 into piggyBac expression vector |
| UL56_Frag2_REV | CTGCAGACGCGTCCAAGGA ATTCTCACCGCCACAGGAAT ACCAG | Subcloning pUL56 AAXA123 into piggyBac expression vector |

**Table S2. TMT sample labelling**

Independent mock infections used in the WT and ΔpUL56 HSV-1 proteomics experiments.

| Experiment | Infection | Time post-infection (h) | Biological replicate | 18-plex TMTpro label |
|---|---|---|---|---|
| WT HSV-1 | WT | 3 | 1 | 126C |
| | WT | 6 | 1 | 127N |
| | WT | 12 | 1 | 127C |
| | WT | 18 | 1 | 128N |
| | WT | 30 | 1 | 128C |
| | WT | 3 | 2 | 129N |
| | WT | 6 | 2 | 129C |
| | WT | 12 | 2 | 130N |
| | WT | 18 | 2 | 130C |
| | WT | 30 | 2 | 131N |
| | Mock | 12 | 1 | 131C |
| | Mock | 12 | 2 | 132N |
| | WT | 3 | 3 | 132C |
| | WT | 6 | 3 | 133N |
| | WT | 12 | 3 | 133C |
| | WT | 18 | 3 | 134N |
| | WT | 30 | 3 | 134C |
| | Mock | 12 | 3 | 135N |
| ΔpUL56 HSV-1 | ΔpUL56 | 3 | 1 | 126C |
| | ΔpUL56 | 6 | 1 | 127N |
| | ΔpUL56 | 12 | 1 | 127C |
| | ΔpUL56 | 18 | 1 | 128N |
| | ΔpUL56 | 30 | 1 | 128C |
| | ΔpUL56 | 3 | 2 | 129N |
| | ΔpUL56 | 6 | 2 | 129C |
| | ΔpUL56 | 12 | 2 | 130N |
| | ΔpUL56 | 18 | 2 | 130C |
| | ΔpUL56 | 30 | 2 | 131N |
| | Mock | 12 | 1 | 131C |
| | Mock | 12 | 2 | 132N |
| | ΔpUL56 | 3 | 3 | 132C |
| | ΔpUL56 | 6 | 3 | 133N |
| | ΔpUL56 | 12 | 3 | 133C |
| | ΔpUL56 | 18 | 3 | 134N |
| | ΔpUL56 | 30 | 3 | 134C |
| | Mock | 12 | 3 | 135N |

**Table S3. TMT-Pro 18-plex reporter ion isotopic impurity values.**

NA, not applicable.

| Mass Tag | Reporter Ion Mass | -2 | | -1 | | M+ | +1 | | +2 | |
|---|---|---|---|---|---|---|---|---|---|---|
| | | -2× 13C | -13C -15N | -13C | -15N | | +15N | +13C | +15N +13C | +2× 13C |
| **TMTpro-126** | 126.127726 | NA | NA | NA | NA | 100% | 0.0034 | 9.31% (127C) | 0.02% | 0.32% |
| **TMTpro-127N** | 127.124761 | NA | NA | NA | 0.78% (126) | 100% | NA | 9.41% (128N) | NA | 0.33% |
| **TMTpro-127C** | 127.131081 | NA | NA | 0.93% (126) | NA | 100% | 0.0035 | 8.63% (128C) | 0.01% | 0.27% |
| **TMTpro-128N** | 128.128116 | NA | 0.00% | 0.95% (127N) | 0.0079 | 100% | NA | 8.38% (129N) | NA | 0.26% |
| **TMTpro-128C** | 128.134436 | 0.00% | NA | 1.47% (127C) | NA | 100% | 0.0034 | 6.91% (129C) | 0 | 0.15% |
| **TMTpro-129N** | 129.131471 | 0.00% | 0.00% | 1.46% (128N) | 0.0128 | 100% | NA | 6.86% (130N) | NA | 0.15% |
| **TMTpro-129C** | 129.13779 | 0.51% | NA | 2.74% (128C) | NA | 100% | 0.0036 | 6.15% (130C) | 0 | 0.11% |
| **TMTpro-130N** | 130.134825 | 0.49% | 0.02% | 2.76% (129N) | 0.0062 | 100% | NA | 5.98% (131N) | NA | 0.11% |
| **TMTpro-130C** | 130.141145 | 0.04% | NA | 3.10% (129C) | NA | 100% | 0.0042 | 4.82% (131C) | 0.02% | 0.06% |
| **TMTpro-131N** | 131.13818 | 0.04% | 0.04% | 3.09% (130N) | 0.0136 | 100% | NA | 4.75% (132N) | NA | 0.06% |
| **TMTpro-131C** | 131.1445 | 0.08% | NA | 3.81% (130C) | NA | 100% | 0.004 | 3.29% (132C) | 0.03% | 0.03% |
| **TMTpro-132N** | 132.141535 | 0.04% | 0.00% | 2.84% (131N) | 0.0079 | 100% | NA | 3.51% (133N) | NA | 0.02% |
| **TMTpro-132C** | 132.147855 | 0.11% | NA | 4.55% (131C) | NA | 100% | 0.0043 | 1.86% (133C) | 0 | 0.00% |
| **TMTpro-133N** | 133.14489 | 0.36% | 0.01% | 3.64% (132N) | 0.0082 | 100% | NA | 1.94% (134N) | NA | 0.00% |
| **TMTpro-133C** | 133.15121 | 0.22% | NA | 4.96% (132C) | NA | 100% | 0.0034 | 1.03% (134C) | 0 | NA |
| **TMTpro-134N** | 134.148245 | 0.40% | 0.04% | 4.92% (133N) | 0.001 | 100% | NA | 1.05% (135N) | NA | NA |
| **TMTpro-134C** | 134.154565 | 0.16% | NA | 5.81% (133C) | NA | 100% | 0.39% | NA (135C) | NA | NA |
| **TMTpro-135N** | 135.151600 | 0.21% | 0.04% | 5.90% (134N) | 0.74% | 100% | NA | NA (136N) | NA | NA |

**Movie S1 (separate file).** Calcium-flux kinetics imaging of i3Neurones mock-infected or synchronously infected (MOI 5) with wild-type or mutant HSV-1. Movie was recorded at 12 hpi and represents 90 s of imaging. 800 μM scale bar.

**Movie S2 (separate file).** Calcium-flux kinetics imaging of i3Neurones mock-infected or synchronously infected (MOI 5) with wild-type or mutant HSV-1. Movie was recorded at 24 hpi and represents 90 s of imaging. 800 μM scale bar.

**Movie S3 (separate file).** Calcium-flux kinetics imaging of i3Neurones mock-infected or synchronously infected (MOI 5) with wild-type or mutant HSV-1. Movie was recorded at 48 hpi and represents 90 s of imaging. 800 μM scale bar.

**Dataset S1 (separate file).** Interactive spreadsheet with "plotter" function to explore multi-omics analysis of HSV-1 infected i3Neurones. "QTV" worksheet enables generation of plots showing changes in gene product abundance by typing in the desired gene name, with proteins and transcripts shown on separate graphs as mean ± SD (three biological replicates) connected by solid and dashed lines, respectively. For each gene product, abundance is normalised from zero to the highest observed value. Data for WT and ΔpUL56 infection are pink and blue, respectively. For proteins, the grey values at time 0 represent the mock infection sample harvested at 12 hpi. For transcripts, the three mock infection time points are each shown (grey). "i3Neurone vs Keratinocyte WCP" worksheet shows the $\log_2$ fold change in whole cell proteomics (WCP) of WT HSV-1 vs mock infection for i3Neurones at 30 hpi and keratinocytes at 18 hpi (2). "PMP" worksheet shows PM proteomics of i3Neurones at 18 hpi. Top graphs show $\log_2$ transformed MaxLFQ abundance of infected with WT (pink) or ΔpUL56 (blue) HSV-1, or mock-infected (grey). Mean ± SD is also shown for three (mock and ΔpUL56 HSV-1) or four (WT HSV-1) biological replicates. Bottom graphs show average $\log_2$ abundances fold change for WT versus mock, ΔpUL56 versus mock or WT versus ΔpUL56 infection.

**Dataset S2 (separate file).** Temporal proteomics analysis of HSV-1 infected i3Neurones. Protein groups (UniProt) and gene names are shown. For worksheets "WT HSV-1 versus Mock" and "ΔpUL56 HSV-1 versus Mock", intensity-Based Absolute Quantification (iBAQ) abundance, number of peptides identified, average $\log_2$ fold change (three biological replicates) of infected cell samples harvested at 3, 6, 12, 18 and 30 hpi versus mock infected samples harvested at 12 hpi, and average normalised reporter ion intensities for each sample are shown. For "WT versus ΔpUL56 HSV-1" worksheet, difference in $\log_2$ fold change versus mock is shown for each time point. For all, FDR-corrected significance (q)-values are shown and significant changes are marked.

**Dataset S3 (separate file).** Temporal transcriptomics analysis of HSV-1 infected i3Neurones. In worksheet "DESqe2 Contrast" the Ensembl gene ID, gene name and Base Mean average normalised count is shown, plus log2 fold change and FDR-adjusted p-value for WT versus mock, ΔpUL56 versus mock and WT versus ΔpUL56 infection at 3, 12 and 30 hpi. "Transcript Counts" worksheet shows the gene ID and normalised counts per transcripts for each replicate experiment at each time point.

**Dataset S4 (separate file).** Plasma membrane proteomics analysis of HSV-1 infected i3Neurones at 18 hpi. "PMP" worksheet shows protein (UniProt) and gene name plus average $\log_2$ fold change (three biological replicates) of WT versus mock, ΔpUL56 versus mock, or WT versus ΔpUL56 HSV-1 infection, with FDR-corrected significance (q)-values are shown and significant changes marked. Average MaxLFQ abundances for each sample are also shown.

**Software S1 (separate file).** R code for filtering and label normalisation of whole cell proteomics data.

**Software S2 (separate file).** R code for DESeq2 analysis of transcriptomics data.

**Software S3 (separate file).** R code for pre-processing of plasma membrane proteomics data.

**Software S4 (separate file).** R code for GO term filtering of plasma membrane proteomics data.

### SI References

1. C. C. Friedel, *et al.*, Dissecting Herpes Simplex Virus 1-Induced Host Shutoff at the RNA Level. *J Virol* **95**, e01399-20 (2021).

2. T. K. Soh, *et al.*, Temporal Proteomic Analysis of Herpes Simplex Virus 1 Infection Reveals Cell-Surface Remodeling via pUL56-Mediated GOPC Degradation. *Cell Rep* **33**, 108235 (2020).

3. S. Tyanova, T. Temu, J. Cox, The MaxQuant computational platform for mass spectrometry-based shotgun proteomics. *Nat Protoc* **11**, 2301–2319 (2016).
