## Supplementary material for "Herpes simplex virus pUL56 abolishes neuronal activity by removing voltage-gated ion channels from the plasma membrane": Software S2

## Load Required Libraries

#Lets load the R and BioConductor packages we need for the analysis.

#```{r}
library(knitr)
library(tximportData)
library(ensembldb)
library(AnnotationHub)
library(tximport)
library(readr)
library(RColorBrewer)
library(Rtsne)
library(gplots)
library(ggplot2)
library(DESeq2)
library(kableExtra)
#```

# Read the sample description information
pdata_full <- read.table(file.path("pdata.txt"), header = TRUE)
rownames(pdata_full) <- pdata_full$technical_sample

pdata <- pdata_full[grep("_1",pdata_full$technical_sample),]
rownames(pdata) <- pdata$sample

# Merge the technical replicate entries to a single entry in a new sample table.
# Populate this with averaged values and point it to the merged quant files in the counts folder.
for (i in 1:nrow(pdata)){
  sample <- pdata[i,"sample"]
  pdata[sample,"raw_reads"] <- sum(pdata_full[pdata_full$sample == sample,"raw_reads"])
  pdata[sample,"mapped_reads"] <- sum(pdata_full[pdata_full$sample == sample,"mapped_reads"])
  pdata[sample,"map_perc"] <- paste0(mean(parse_number(pdata_full[pdata_full$sample == sample,"map_perc"])),"%")
  pdata[sample,"run"] <- paste(as.character(pdata_full[pdata_full$sample == sample,"run"]),collapse="_")
  pdata[sample,"filename"] <- paste0("counts/fastq_merged/",sample,"_quant/quant.sf.gz")
}

pdata <- pdata[order(as.numeric(gsub("t","",pdata$time)),pdata$treatment),]

# Print out the main sample table
kable(pdata,label = "Sample Description")


# Check the quantification files from salmon exist
files <- file.path(paste0("./counts/",pdata_full$filename))
names(files) <- pdata_full$technical_sample

files_merged <- file.path(pdata$filename)
names(files_merged) <- pdata$sample

if(all(file.exists(files),file.exists(files_merged))){
  print("[Passed] All quant files listed exist")
} else {
  print("[Error ] Some quant files are missing...")
}


#```
## Colour Palettes

#Let's set up colour palettes for heatmaps and plots later on. We will make unique schemes for sample type, time and treatment.
#```{r fig.retina = 2, dpi=300, fig.height= 6 }

show_pal <- function(palette,names=NULL,main=NULL,cex=NULL) 
{
  n <- length(palette)
  if (length(palette > 0)) {
    image(1:n, 1, as.matrix(1:n), col = palette, xlab = "", 
          ylab = "", xaxt = "n", yaxt = "n", bty = "n")
    text(1:n, 1, names,cex=cex)
  }
  if(!is.null(main)){
    title(main)
  }
}
# Heatmap color scheme
hmcol = colorRampPalette(brewer.pal(9, "GnBu"))(100)

# Make a colour scheme for sample type

type_col = colorRampPalette(brewer.pal(9, "Set1"))(length(unique(pdata$type)))
names(type_col)=unique(pdata$type)

time_col = colorRampPalette(brewer.pal(9, "BuPu"))(length(unique(pdata$time)))
names(time_col)=unique(pdata$time)

treat_col = colorRampPalette(brewer.pal(3, "Paired"))(length(unique(pdata$treatment)))
names(treat_col)=unique(pdata$treatment)

par(mfrow=c(4,1))
show_pal(hmcol,main="Heatmap")
show_pal(type_col,cex=0.6,names=names(type_col),main="Type")
show_pal(time_col,cex=0.6,names=names(time_col),main="Time")
show_pal(treat_col,cex=0.6,names=names(treat_col),main="Treatment")
par(mfrow=c(1,1))
#```


## Load Human Annotation Data

#We will use the `AnnotationHub` package and `ensembldb` to load the v113 Ensembl transcript data into R. We'll use this to process the salmon quant files individually and allow it to merge counts from multiple transcripts back to the gene-level.

#Because we've engineered in HSV1 genes, we'll do a little hack such that the 77 HSV transcripts are also processed.


#```{r}
# Activate Annotation Hub
ah <- AnnotationHub()

# If you need to see available annotations, this command can be used, we will choose AH119325
#query(ah, "EnsDb.Hsapiens") 

# The one we want is: 
# AH119325 | Ensembl 113 EnsDb for Homo sapiens
#

# Make Ensembl Database for that human genome annotation version
edb <- ah[["AH119325"]]

# Build a tx2gene table for the human transcripts, discard un-needed entries, but keep full table if useful for annotating things later 
tx2gene <- transcripts(edb, return.type = "DataFrame")
tx2gene$tx_external_name <- gsub("-\\d+","",tx2gene$tx_external_name)
tx2gene_simple <- tx2gene[,c(1,7,11,2)]

# Read in the HSV specific annotation table
hsv <- read.table("genome/hsv_gene_table.txt",header=FALSE,sep="\t")

# We'll add all 77 HSV cDNAs onto the end of our tx2gene_simple table
for(i in 1:nrow(hsv)){
  tx2gene_simple[nrow(tx2gene_simple) + 1,] = c(hsv[i,c(1,2)],hsv[i,2],"viral")
}

# Lets import the salmon quantifications using tximport, Ensembl version numbers caused issues, so ignore them
txi.salmon.full <- tximport(files, type = "salmon", tx2gene = tx2gene_simple,ignoreTxVersion = TRUE)
txi.salmon <- tximport(files_merged, type = "salmon", tx2gene = tx2gene_simple,ignoreTxVersion = TRUE)

kable(head(txi.salmon$counts))


## Technical Replication and Merging

#Libraries were split across two flow-cells. Let's explore the correlation of samples between those technical replicates. The Heatmap below seems to show excellent correlation between the technical replicates as expected and no libraries appear to have dropped out.

#```{r fig_width=7, fig_height=10, dpi=300}
heatmap.2(cor(txi.salmon.full$counts),col=hmcol,trace="none",cexCol = 0.4,cexRow = 0.4,main="Technical Replicate Correlation")
#```


gene_names=tx2gene_simple
rownames(gene_names)=gene_names$gene_id


dds <- DESeqDataSetFromTximport(txi.salmon, pdata, ~type)


# Filter the data perhaps for low abundance reads and viral reads - may improve the fit
#dds <- dds[grep("^hsv1_",rownames(counts(dds)),invert = TRUE),]
dds <- dds[apply(counts(dds),1,median) > 3,]
dds <- estimateSizeFactors(dds)
dds <- estimateDispersions(dds)
plotDispEsts(dds)
dds = nbinomWaldTest(dds)


contrasts <- list(
  c("wt_t3", "cmock_t3"),
  c("wt_t12", "cmock_t12"),
  c("wt_t30", "cmock_t30"),
  c("d56_t3", "wt_t3"),
  c("d56_t12", "wt_t12"),
  c("d56_t30", "wt_t30"),
  c("d56_t3", "cmock_t3"),
  c("d56_t12", "cmock_t12"),
  c("d56_t30", "cmock_t30")
)


res_list <- list()
contrast_names <- c()

# Loop over all your contrasts
for (i in 1:length(contrasts)) {
  c1 <- contrasts[[i]][1]
  c2 <- contrasts[[i]][2]
  contrast_name <- paste0(c1, "_vs_", c2)
  contrast_names <- c(contrast_names, contrast_name)
  
  res <- results(dds, contrast=c('type', c1, c2))
  colnames(res) <- paste0(colnames(res), "_", contrast_name)
  res_list[[i]] <- as.data.frame(res)
}

# Before merging, add gene_id column to each data.frame
res_list_named <- lapply(res_list, function(df) {
  df$gene_id <- rownames(df)
  return(df)
})

# Merge safely on gene_id
full_table <- Reduce(function(x, y) merge(x, y, by = "gene_id", all = TRUE), res_list_named)

# Set gene_id as rownames
rownames(full_table) <- full_table$gene_id
full_table$gene_name <- gene_names[rownames(full_table), "tx_external_name"]
full_table <- full_table[, c("gene_id", "gene_name", setdiff(names(full_table), c("gene_id", "gene_name")))]

# Add in normalized counts:
# Step 1: Get normalized counts
norm_counts <- as.data.frame(counts(dds, normalized = TRUE))


# Use the 'type' from pdata to relabel
colnames(norm_counts) <- make.unique(pdata[colnames(norm_counts), "type"])

# Step 2: Add gene_id as a column for merging
norm_counts$gene_id <- rownames(norm_counts)

# Step 3: Merge with full_table by gene_id
final_table <- merge(full_table, norm_counts, by = "gene_id", all.x = TRUE)


write.table(final_table,
            file = "251212_DESeq2_all_contrasts_withvirus_results.txt",
            sep = "\t",
            quote = FALSE,
            row.names = FALSE)
